# Over-expression of the photoperiod response regulator *ZmCCT10* modifies plant architecture, flowering time and inflorescence morphology in maize

**DOI:** 10.1101/402586

**Authors:** Elizabeth Stephenson, Stacey Estrada, Xin Meng, Jesse Ourada, Michael G. Muszynski, Jeffrey E. Habben, Olga N. Danilevskaya

**Affiliations:** CORTEVA Agrisciences, Agriculture Division of DowDuPont; Johnston, Iowa, United States of America; University of Hawaii at Manoa, Tropical Plant and Soil Sciences, Honolulu, Hawaii; United States of America

## Abstract

Maize originated as a tropical plant that required short days to transition from vegetative to reproductive development. *ZmCCT10* [CO, CONSTANS*, CO-LIKE* and *TIMING OF CAB1* (*CCT*) transcription factor family] is a photoperiod regulator and was identified as a major QTL controlling photoperiod sensitivity in maize. We modulated expression of *ZmCCT10* in transgenic maize using two constitutive promoters which cause differing expression levels. Transgenic plants over expressing *ZmCCT10* with either promoter was delayed in their transition from vegetative to reproductive development but were not affected in their change from juvenile-to-adult vegetative growth. Strikingly, transgenic plants containing the stronger expressing construct had a very prolonged period of vegetative growth accompanied with dramatic modifications to plant architecture that impacted both vegetative and reproductive traits. These plants did not produced ears, but tassels were heavily branched, and more than half of the transgenic plants showed conversion of shoot apices into “bushy tops”, which were composed of vegetative reversion plantlets. Analysis of expression modules controlling the floral transition and meristem identity linked these networks to photoperiod dependent regulation, whereas phase change modules appeared to be photoperiod independent. Results from this study clarified the influence of the photoperiod pathway on vegetative and reproductive development and allowed to fine-tune the flowering time model for maize.

## Introduction

Plants display an astonishing diversity in body plan architecture [1]. However much of-the complexity of plant morphology can be explained by variations of the basic architectural unit – the phytomer. The phytomer is comprised of the leaf, attached to the node on the stem, the internode, the stem segment between two successive nodes, and the axillary bud positioned in the axil of the leaf [1]. The number of phytomers is determined by the activity of the shoot apical meristem (SAM), or the apical bud, which is a pool of undifferentiated pluripotent stem cells capable of producing above ground organs [2]. The axillary bud is also composed of meristematic cells which initiate the growth of side branches [3]. The SAM represses the outgrowth of the axillary bud to control the extent of branching which is known as apical dominance [4].

Maize is an annual grass with a determinant habit of growth. The main stalk of the mature plant is composed of a variable number of phytomers depending on genotype. The apex of the plant terminates when the SAM converts into the male inflorescence, called the tassel. The female inflorescence, the ear, is initiated from axillary buds. These buds form in leaf axils but usually only the top one or two buds on the plant develop into ears and bear seed; whereas the ears positioned lower on the shoot abort [5].

After germination, maize seedlings transition through distinct developmental phases. Vegetative development is divided into two phases - juvenile and adult [6]. Seedlings in the juvenile phase display characteristic traits such as the presence of adventitious roots, short internodes, and narrow leaves [7]. Juvenile leaves also have a number of distinctive epidermal traits including weakly invaginated cell walls, epicuticular waxes, the absence of trichomes (macrohairs) and the presence of bulliform cells [7-9]. The transition from the juvenile to adult phase is regulated by the balance of two micro-RNAs, miR156 and miR172, where miR156 promotes the juvenile phase and miR172 promotes the adult phase [10, 11].

High expression of miRNA156 maintains the juvenile phase by repression of *SQUAMOSA PROMOTER BINDING PROTEIN (SBP/SPL)* genes, which encode plant-specific transcription factors required for development of adult traits [11, 12]. In maize, over-expression of miR156 results in the striking grass-like phenotype of the *Corngrass1* (*Cg1*) mutant [12]. This mutant retains juvenile traits such as short internodes, multiple tillers, slender culms, an increased number of vegetative nodes, adventitious roots, narrow leaves with epicuticular wax and the absence of trichomes [13]. Transcripts of at least seven *SBP/SPL* genes are down regulated in the *Cg1* mutant including *teosinte glume architecture1* (*tga1*), a *SBP* transcription factor involved in maize domestication [12]. Upon seedling growth expression of miR156 declines allowing expression of *SBP* genes to increase, leading to the transition of adult growth [6, 10, 11].

By contrast, miR172 functions antagonistically to miR156 by repressing juvenile traits and accelerating the transition to the adult phase by down regulation of *APETALA2*-like (*AP2*) transcription factor genes [10, 11]. The key regulator of the juvenile-to-adult transition in maize is the AP2-like *GL15* (*glossy 15*) gene [7, 14]. Over-expression of miR172 degrades *GL15* mRNA that results in an accelerated transition to the adult phase [15]. Genetically, *glossy15* functions downstream of *Cg1* [7, 14], which is consistent with molecular data showing expression of miR172 transcripts are reduced in *Cg1* mutants [12].

After the juvenile-to-adult phase change, plants acquire the competence for reproductive development. The switch from adult vegetative growth to reproductive growth is called floral transition. During floral transition the SAM ceases leaf initiation and is transformed into the inflorescence meristem (IM). In maize, this is marked by the SAM becoming committed to tassel development [16]. The total number of leaves produced is often used as a quantitative measurement of the length of the vegetative stage of growth. The transition to reproductive development is regulated by numerous environmental and endogenous cues that stimulate accumulation of the flowering hormone florigen in leaves. Florigen is transmitted from leaves to the shoot apical meristem through the phloem where the transition to reproductive development occurs [17-20].

Florigen was identified in Arabidopsis as the 23 kD protein encoded by the *FLOWERING locus T (FT)* gene [17, 21]. Afterwards, homologs of the *FT* gene were detected in virtually all plants and the FT protein is postulated to be the universal flowering hormone [18, 22, 23]. In the SAM, FT interacts with a 14-3-3 receptor protein and the bZIP transcription factor FLOWERING LOCUS D (FD) forming the florigen activation complex (FAC) as demonstrated in rice [24, 25]. The FAC activates transcription of the *APETELA 1 (AP1)-like MADS* box genes which marks the onset of reproductive development [24]. Formation of the FAC seems to be a universal feature in flowering plants but direct evidence of a FAC beyond rice is lacking [26].

The *FT-FD* genetic module is conserved in maize. The *delayed flowering1* (*dlf1*) gene encodes an *FD*-like bZIP transcription factor which mediates floral signals in the shoot apex [27]. The expanded family of *FT*-like genes in maize were named *Zea CENTRORADIALIS* (*ZCN*) reflecting their functional diversification [28]. A florigenic function was shown for the *ZCN8* gene [29, 30]. Its nearly identical paralog *ZCN7* is thought to also possess florigenic activity [31]. Flowering Zn-finger transcription factor *indeterminate1* (*id1*)[32] controls expression of *ZCN8* and *ZCN7* [29, 31]. Although it is not clear how *id1* regulates *ZCN7* and *ZCN8*, recent data suggests it may be via epigenetic modification of their chromatin structure [31]. It is likely that the FAC is also formed in maize because the ZCN8 protein was shown to interact with the DLF 1 protein [30]. Similar to Arabidopsis and rice, the onset of reproductive development in maize is demarcated by expression of the *AP1*-like MADS box genes *ZMM4* and *ZMM15* [33], which supports a conserved developmental genetic pathway between these species.

Maize was domesticated from the tropical grass teosinte (*Zea mays* ssp. *parviglumis*) that requires short days to flower [34]. Over time as maize cultivation moved to higher latitudes with longer summer days, selection for short day sensitivity was significantly weakened [35]. There is wide variation in the photoperiod sensitivity among maize inbred lines ranging from complete day length insensitivity, to moderately and highly sensitive groups [36]. Day neutral lines produce equal number of leaves under both short (SD) and long (LD) days. Gaspé Flint, which is the earliest known cultivar of maize, produces on average 10 leaves under both conditions. The highly day-length sensitive tropical lines can produce up to 30 leaves under LDs compared to 23 leaves under SDs [36], which indicates that tropical maize and teosinte are facultative SD plants because they still flower under non-inductive LDs; while inductive SDs accelerate flowering.

Using a maize-teosinte mapping population and genome-wide association the major photoperiod response regulator, the *ZmCCT10* gene, was identified on chromosome 10 [37]. The *ZmCCT10* gene encodes a CCT (CO, CO-LIKE and TIMING OF CAB1) domain protein. Further studies revealed the insertion of a CACTA-like transposon in the upstream promoter regions of the *ZmCCT10* gene in day-neutral temperate lines which presumably disrupts *ZmCCT10* expression and attenuates photoperiod sensitivity under LD conditions [38]. Recently the *ZmCCT9* gene, on chromosome 9, was also identified to control flowering similar to *ZmCCT10* [39].

The maize *ZmCCT10* is a homolog of the rice photoperiod response regulator *Ghd7*, which was identified as a quantitative trait locus (QTL) for Grains, plant height and heading date on chromosome 7 [37, 40]. *Ghd7* is a negative regulator of flowering (heading date) [40, 41] and is expressed in leaves with a diurnal pattern, peaking in the early morning under LDs. In contrast, its expression is low under SD [40]. The GHD7 protein represses transcription of the floral inducer *Early Heading Date1 (Ehd1*) which promotes flowering under SDs [42]. EHD1 is a B-type response regulator with DNA binding properties suggesting it functions as a transcription factor [42]. *Ehd1* is required for expression of the rice florigen gene *Heading Date3a* (*Hd3a)* [23] and *Rice flowering locus T1* (*RTF1*), major floral activators under LD conditions [43, 44]. The floral promoter *Ehd1* and the floral repressor *Ghd7* fine tune the expression of the *Hd3a/RFT1* genes to recognize a critical day length for transition to reproductive development [45].

The *Ghd7-Ehd1* genetic module seems to be conserved in other short day tropical crops as was demonstrated for sorghum [46, 47] and also for maize [38, 48]. In the long day winter crops wheat and barley, *VERNILIZATION2* (*VRN2*), a homolog of *Ghd7,* represses *FT*-like genes prior to cold exposure [49, 50]. But the *Ghd7-Ehd1* module is not present in Arabidopsis or other eudicots [40, 51].

*Ghd7* over-expression or down-regulation in transgenic plants revealed its role as a central regulator of growth, development and stress response in rice [41]. Over-expression of *Ghd7* affected plant architecture resulting in taller plants with thick stems, fewer tillers but increased panicle branching which led to more grain per plant [40, 41]. *Ghd7* is one of the major targets for increasing grain yield in rice breeding programs [52].

Our knowledge of how the *ZmCCT10* gene functions at the molecular level is limited [38]. However, being a repressor of the photoperiod pathway, manipulation of *ZmCCT10* expression provides the opportunity to study the role of this pathway in floral transition and other developmental processes. To investigate these roles, we over-expressed *ZmCCT10* driven by constitutive promoters of different strengths in the day neutral early-flowering maize line Gaspé Flint. The resulting transgenic plants displayed dramatic modification of plant morphology producing tall, late flowering phenotypes with about 50% of the events showing vegetative reversions in the tassel producing a “bushy top” phenotype.

## Methods

### Plant materials

Extremely early temperate cultivar Gaspé Flint, temperate inbred line B73, tropical CML436 and CML311maize lines were used due to their distinct photoperiod sensitivities and differences in flowering times. Diurnal experiment using these lines was described in [30]. Teosinte lines were obtained from North Central Regional Plant Introduction Station, Ames, Iowa, USA. Teosinte accession# PI 441934 is *Zea mays* sp *huehuetenangensis* originally from Guatemala. Teosinte accession # PI 422162 is *Zea luxurians* originally from Mexico.

### T-DNA constructs and plant transformation

GATEWAY®TECHNOLGY (Invitrogen, CA) was used for vector construction. The co-integrated vectors were constructed, and plants were transformed as described previously [30, 53] into Gaspé Flint. Typically, 10 independent single copy events were generated for each construct. T1 seeds were generated by pollination with Gaspé Flint as a pollen donor.

### Phenotypic data collection

Vegetative growth stages (V stage) were defined according to the appearance of the leaf collar of the uppermost leaf [54]. The staging notes were taken twice a week. Using staging notes leaf appearance rate was calculated with linear regression models. Growth rate calculated as (H_F_-H_1_)/(DAP_F_-DAP_1_) where H -height, F - final height, 1 - first recorded measurement, where DAP stands for days after planting. Leaf length was measured on fully expanded leaves with a visible ligule as the distance from the leaf collar/ligule to the tip of the blade. Leaf width was measured at the widest point of the blade. The ratio of the length to the width was calculated. Nodes were identified by the leaf number originating from that node. Internode lengths were measured when fully mature plants were harvested and were calculated as the distance between nodes. Only above-ground internodes were measured.

### Toluidine blue O staining of epidermal peels

To determine juvenile to adult phase transition sections from leaf 2 and up until adult traits were confirmed were sampled. Leaf segments from margin to margin were collected from the base and the tip of a single leaf. Segments were fixed in a mixture of 1-part ethanol to 3 parts acetic acid [55]. The abaxial epidermis and mesophyll were removed using abrasive techniques. Once cleared, the adaxial epidermis was stained with 0.05% toluidine blue O/acetic acid, pH 4.5 (TBO solution) [7, 56] for 30 seconds. The peel was then rinsed with deionized water and immediately photographed using bright-field optics on a microscope.

### Tissue collection for qRT-PCR

Plants were grown in the greenhouse under LD conditions with a 14-hour day length. To cover much of the developmental range of candidate gene expression, coleoptiles were sampled from NTG and UBI_pro_:*ZmCCT10* transgenic plants but not from the BSV_pro_:*ZmCCT10* plants due to limited seed availability. Afterwards, the first true leaf was sampled, and then leaves were sampled (with a punch) twice a week beginning from leaf # 4. Due to expression of the maize florigen gene *ZCN8* at the tip of the leaf [30], leaves were consistently sampled 3-5 cm from the tip. To capture diurnal expression patterns, leaves were sampled in the morning when lights were turned on in the greenhouse and 12 hours later in the evening. When the BSV_pro_:*ZmCCT10* plants reached 24 leaves, it was technically challenging to sample such tall plants and thus leaves 24-31 were sampled at the final dissection of the plants in the afternoon. Tissues collected for qRT-PCR were immediately flash frozen after being sampled. Additional NTG and UBI_pro_:*ZmCCT10* transgenic plants were grown in a Conviron CMP6050 growth chamber under 14-hour days (28°C days and 26°C nights) to sample meristems. Meristems were dissected at every V stage starting at V1 until the meristem transitioned to a tassel (∼V4 for NTG and ∼V4 to V7 for UBI_pro_:*ZmCCT10* transgenic plants depending on the CCT10 allele). Due to limited seed availability, BSV_pro_:*ZmCCT10* plant meristems were sampled in the greenhouse under LD conditions with a 14-hour day length during the initial T1 experiment. Plants were sampled at various stages ranging from V6 to V32. Images were taken of each meristem with a Leica MSV269 dissecting scope, then immediately flash frozen to be used for qRT-PCR.

### RNA isolation and qRT-PCR

Total RNA was isolated using Qiagen RNeasy reagents (https://www.qiagen.com/us/shop/Lab-Basics/Buffers-and-Reagents/) with the nucleic acid bound to columns of a 384 well binding plate purchased from the Pall Corporation (http://www.pall.com/main/oem-materials-and-devices/product.page?lid=gri78l6g). DNA was removed from the RNA samples using Roche DNAse I Recombinant (https://www.lifescience.roche.com/shop/en/us/products/dnase-i-recombinant-rnase-free) and synthesis of cDNA was done using Applied Biosystems High Capacity cDNA Reverse Transcription kits (https://www.thermofisher.com/order/catalog/product/4368813?ICID=search-product). Quantitative PCR was done using hydrolysis probe and SYBR based reactions. Primers and probes were designed using Applied Biosystems Primer Express software (https://www.thermofisher.com/order/catalog/product/4363993?ICID=search-product) using nucleotide sequences published in Genbank. Hydrolysis probe-based PCR was performed using Bioline Sensi-fast mix (http://www.bioline.com/us/sensifast-probe-lo-rox-kit.html) while SYBR-based PCR was run using Applied Biosystems PowerUp SYBR Green Master Mix (https://www.thermofisher.com/order/catalog/product/A25741?ICID=search-product). All reactions were run on an Applied Biosystem Viia7 Real-Time PCR instrument using the manufacturer’s conditions. Relative gene expression was calculated by normalizing against *maize eukaryotic initiation factor 4-gamma* gene (GenBank accession # EU967723). A list of primers and probes are shown in S5 Table. To identify genes which expressions were different in transgenic apices compared to NTG plants, T-test was performed for every developmental stage. Expression level were considered statistically significant with p<0.05 (S6 Table).

## Results

### Over-expression of *ZmCCT10* produces dramatic effects on multiple traits in T0 maize plants

To investigate how over expression of *ZmCCT10* may impact flowering, we used a transgenic approach and chose to constitutively over-express the *ZmCCT10* coding region. To explore how diverse allelic variation of the ZmCCT10 protein might differentially alter flowering, *ZmCCT10* alleles from different maize inbred lines with distinct flowering characteristics were selected, including the day-length neutral early flowering Gaspé Flint, the temperate B73, and the SD-sensitive tropical CML436 and CML311 lines [30]. Two teosinte accessions, the SD-sensitive wild progenitor of maize, were also used (accessions PI 441934 and PI 422162). To identify conserved protein domains, maize ZmCCT10 and sorghum SbGHD7, the closest maize CCT10 homolog which functions as a floral repressor under long days, were compared [46]. Their amino acid alignment showed the proteins were conserved with 61% amino acid identity (S1 Fig).

To investigate how the level of *ZmCCT10* expression could affect flowering, we used two constitutive promoters of different strengths to drive expression of six different alleles. The maize ubiquitin promoter (UBI_pro_) is widely used in cereal transgenic studies as a constitutive promoter with a high level of expression [57]. However, the Banana Streak Virus promoter (BSV_pro_) exceeds the level of expression driven by the UBI_pro_ and directs expression in all tissue tested with exception of pollen [58]. Using both the UBI_pro_ and BSV_pro_ and the genomic and/or cDNA sequences of six *ZmCCT10* alleles, we constructed a cohort of constructs that provided a wide range of allelic and expression combinations in which to study the effect of *ZmCCT10* overexpression on plant phenotype (S1 Table).

All constructs were transformed into the day-neutral early flowering Gaspé Flint line (Fig 1A). Between 9-10 single copy transgenic events were generated for each construct. Because T0 transgenic plants were produced in batches, representative T0 data were collected for each experimental batch grown in the greenhouse at the same time (S1 Table).

**Fig 1.**
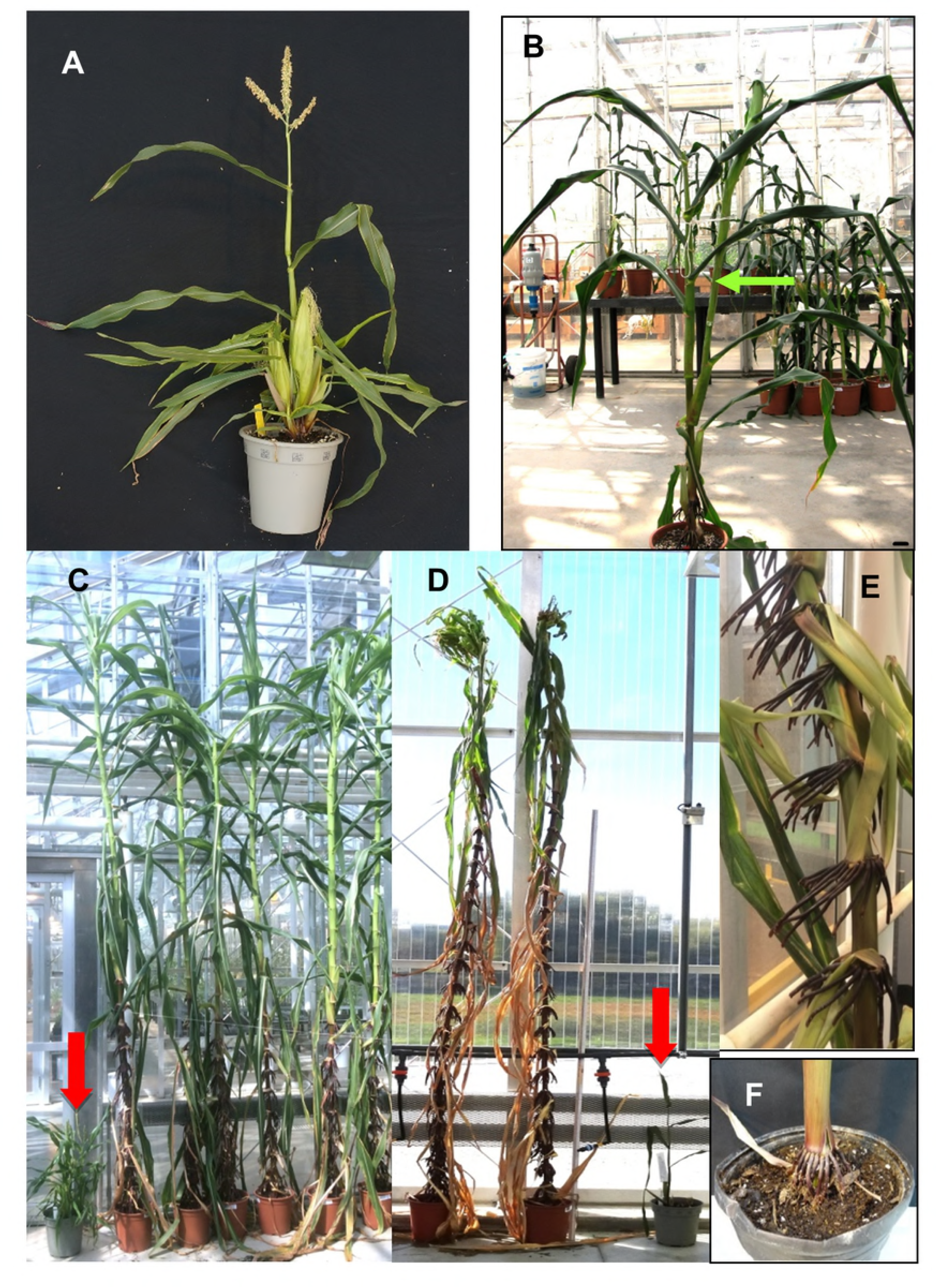
Representative images of non-transgenic (NTG), UBIpro:*ZmCCT10*, and BSVpro:*ZmCCT10* transgenic plants. A) Image of a non-transgenic (NTG) Gaspé Flint line used for transformation. B) UBIpro:*ZmCCT10* transgenic plant with a long ear shank (green arrow). C) BSVpro:*ZmCCT10* transgenic plants at 119 days after planting and a Gaspé Flint parental line (red arrow). D) BSVpro:*ZmCCT10* transgenic plants at 171 days after planting and a Gaspé Flint parental line (red arrow). Note unusual bushy structures are formed at the top of transgenic plants. E) A closeup view of extensive brace roots on a BSVpro:*ZmCCT10* transgenic plant at the 15-20 internode region. F) A typical Gaspé Flint plant with brace roots only at the base of the plant.

As expected for over-expression of a flowering repressor, the transgenic plants exhibited a late flowering phenotype which was manifested as taller plants with more leaves compared to the non-transgenic (NTG) parental Gaspé Flint line (Fig 1B and 1C). Delayed flowering, recorded as time to pollen shed (shedding) and silk exertion (silking), was also observed (S1 Table). Typical time for Gaspe Flint to shed and silk is about 30 days.

In addition to the expected late flowering phenotype, the transgenic plants also displayed unexpected features. The UBI_pro_:*ZmCCT10* T0 plants produced atypically long axillary branches subtending the ears (Fig 1B). Moreover, BSV_pro_:*ZmCCT10* transgenic plants exhibited a severe modification to the entire plant architecture (Fig 1C and 1D). The BSV_pro_:*ZmCCT10* transgenic plants were extremely tall, up to 300 cm, on average, and produced up to 38 leaves (Fig 1C, 1D and S2A Fig; S1 Table). The stalks of BSV_pro_:*ZmCCT10* transgenic plants were strong yet flexible with increased mechanical strength and could withstand bending up to 80° (S3 Fig). Brace roots formed up to the 35^th^-37^th^ nodes compared to the base internode of NTG plants (Fig 1E and 1F, S2B Fig). Secondary aerial brace roots developed as the plants aged (S2D Fig). In contrast, the non-transgenic Gaspé Flint line produced only a few brace roots at the base of the plant (Fig 1F). Unexpectedly, as plants matured, a vegetative, highly branched structure emerged from the top of many of the BSV_pro_:*ZmCCT10* plants in lieu of tassels (Fig 1C). More than 50% of the BSV_pro_:*ZmCCT10* transgenic plants produced these vegetative apical structures that we called “bushy top” (S2E Fig) and were composed of reversion plantlets. This phenotype was not observed in any of the NTG or UBI_pro_:*ZmCCT10* plants.

### Pleiotropic effects of *ZmCCT10* are independent of allelic variations but are dosage dependent

We further examined *ZmCCT10* overexpression effects in the T1 families. We selected 12 constructs composed of six cDNA *ZmCCT10* alleles driven by either the UBI_pro_ or BSV_pro_ (S2 Table). The T1 families of 20 plants from two events were planted for the UBI_pro_:*ZmCCT10* constructs. Due to the reduced fertility of the BSV_pro_:*ZmCCT10* transgenic plants, the available T1 seed was limited and, in a few cases, only 3-4 plants were planted (S2 Table). Consistent with T0 observations, both UBI_pro_:*ZmCCT10* and BSV_pro_:*ZmCCT10* constructs modified plant morphology in the T1 plants, with the more extreme manifestations being observed in the BSV_pro_:*ZmCCT10* plants. A weak allelic effect was detected in the UBI_pro_:*ZmCCT10* constructs where UBI_pro_:*ZmCCT10*_Gaspé_ and UBI_pro_:*ZmCCT10*_B73_ showed a smaller effect on plant height and leaf number. The allelic effect was not observed in the BSV_pro_:*ZmCCT10* plants and we surmise that this lack of phenotypic effect may have been overridden by the very high level of transgene expression (S2 Table). Thus, we considered the level of transgene expression as the major factor correlated with severity of phenotype. For this reason, we analyzed data by grouping the UBI_pro_:*ZmCCT10* and BSV_pro_:*ZmCCT10* results regardless of their allele background.

The T1 families showed a strong association between the level of transgene expression and trait measured. This occurred for plant height (S4A Fig), leaf number (S4B Fig), the uppermost node with brace roots (S4C Fig), and the primary ear position (S4D Fig). Higher levels of transgene expression resulted in enhancement of the trait modification. However, there was a threshold for transgene expression in the BSV_pro_:*ZmCCT10* plants beyond which trait enhancement plateaued (S4 Fig).

### Modification of vegetative traits in T1 maize transgenic plants

The T1 transgenic families exhibited modification of plant phenotypes consistent with the T0 generation. The average BSV_pro_:*ZmCCT10* T1 plant height was 263 cm compared to 105 cm for the UBI_pro_:*ZmCCT10* plants and ∼ 70 cm for the NTG siblings (Table 1). The growth rate of the BSV_pro_:*ZmCCT10* plants was only 2.8 cm/day compared to ∼ 4.2 cm/day of the UBI_pro_:*ZmCCT10* and NTG plants (Table 1). Despite their slower growth rate, the BSV_pro_:*ZmCCT10* transgenic plants produced more internodes because they remained in a vegetative stage longer, which also resulted in the increased plant height.

**Table 1.** Vegetative traits collected for T1 families 1.

| Line | Total No. Plants | Relative Expression of *ZmCCT10* qRT-PCR | Plant Height (cm) | Growth rate (cm/day) | Leaf No. | Leaf appearance rate (leaves/day) | Top Nodes with brace roots | Nodes with initiated primary ears |
| --- | --- | --- | --- | --- | --- | --- | --- | --- |
| NTG siblings | 60 | 0 | 74.4 ± 14.2 | 4.4 ± 1.0 | 9.8 ± 0.8 | 0.28 | 4 to 6 | 6 to 7 |
| UBI<sub>pro</sub>:*ZmCCT10* | 120 | 2.7 ± 1.3 | 104.9 ± 29.8* | 4.3 ± 0.8 | 12.6 ± 2.1* | 0.30 | 5 to 9 | 6 to 8 |
| NTG siblings | 45 | 0 | 67.9 ± 16.2 | 4.1 ± 1.1 | 9.5 ± 0.8 | 0.28 | 4 to 6 | 6 to 7 |
| BSV<sub>pro</sub>:*ZmCCT10* | 69 | 11.9 ± 5.7 | 263.0 ± 51.8** | 2.8 ± 0.7 | 35.9 ± 5.8** | 0.31 | 16 to 37 | 18 to 33 |

Measurements represent means ± SD. Leaf appearance rate calculated with linear regression models. BSV<sub>pro</sub>:*ZmCCT* ear traits were collected only for one event (11 plants). \*Means are statistically different from NTG at p < 0.001. \*\*Means are statistically different from NTG at p < 0.00001.

Internode length depends on position within the plant shoot. As observed for NTG and UBI_pro_:*ZmCCT10* plants, the internodes below the ear were progressively longer closer to the ear and then became shorter above the ear (Fig 2A). Thus, internode length displayed an acropetal, bottom-up gradient below the ear and then switched to a top-down basipetal gradient above the ear, which is characteristic for organ growth before and after floral transition [4]. The BSV_pro_:*ZmCCT10* internodes were on average 1.5-fold shorter and 2-fold wider than NTG and UBI_pro_:*ZmCCT10* plants (Fig 2A and S5 Fig). Diameters of the BSV_pro_:*ZmCCT10* stalks measured at the 1^st^ internode averaged 31 mm compared to 15 mm for NTG and 20 mm for UBI_pro_:*ZmCCT10* plants (S3 Table). This observation may explain the slower relative growth of the BSV_pro_:*ZmCCT10* plants since they are allocating more assimilates to stalk-width growth.

**Fig 2.**
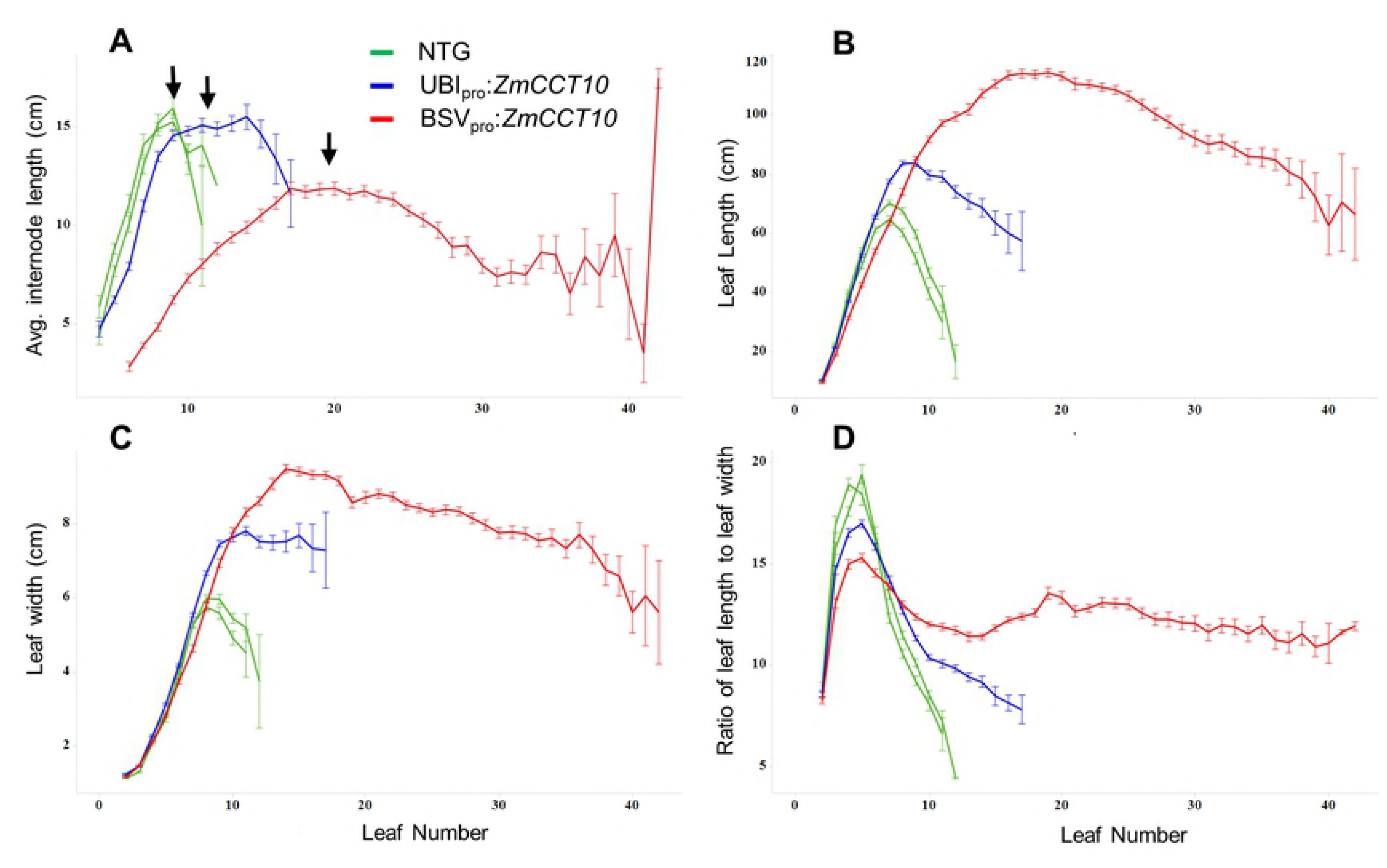
Internodes length and leaf morphology in non-transgenic (NTG), UBIpro:*ZmCCT10*, and BSVpro:*ZmCCT10* transgenic plants. A) Internode length by node position, B) leaf length by leaf position, C) leaf width by leaf position, and D) length/width ratios by leaf position. Node and leaf number are numbered from the base to the apex of the shoot. The ear nodes marked by black arrows. Error bars represent the ± the SE.

The UBI_pro_:*ZmCCT10* plants produced an average of 13 leaves compared to 9-10 leaves for NTG plants, whereas the BSV_pro_:*ZmCCT10* plants produced 36 leaves (Table 1). Leaf appearance rate was accelerated in transgenic plants: 0.30 and 0.31 leaves/day in UBI_pro_:*ZmCCT10* and BSV_pro_:*ZmCCT10* plants respectively versus 0.28 leaves/day in NTG plants (Table 1 and S6 Fig). This observation suggested that leaves in the SAM were initiated at a faster rate in transgenic plants relative to NTG plants.

Leaf size showed an acropetal growth pattern below the ear and a basipetal pattern above the ear (Fig. 2). In NTG and UBI_pro_:*ZmCCT10* plants, leaves below the ear displayed a sharp increase in length and width closer to the ear resulting in an increased length/width ratio. Leaves above the ear became shorter and narrower and the length/width ratio sharply declined (Fig 2B, 2C and 2D). Typically, the biggest leaf on a plant is the ear leaf. The BSV_pro_:*ZmCCT10* plants displayed the characteristic acropetal pattern of leaf size but the basipetal pattern was disrupted after the 10^th^ leaf. Leaf size remained relatively constant above node 10 reflecting the longer and wider leaf shape (Fig 2B, 2C, and 2D). This indicated an increased leaf growth rate, longer duration of growth, or both in the BSV_pro_:*ZmCCT10* plants compared to NTG or UBI_pro_:*ZmCCT10* plants.

Consistent with the T0 observations, brace roots were found as high as node 37 in the BSV_pro_:*ZmCCT10* plants, which was above the node of primary ear formation (Table 1); whereas NTG and UBI_pro_:*ZmCCT10* plants produced brace roots below the ear (NTG internode 4-6 and UBI_pro_:*ZmCCT10* internode 5-9). Because brace roots typically form at juvenile nodes [7] we decided to examine the phase change in T1 families. To determine juvenile and adult growth phases we first observed the presence of macro-hairs (an adult trait). Macro-hairs were present in NTG, UBI_pro_:*ZmCCT10* and BSV_pro_:*ZmCCT10* plants starting at leaf #4 (S4 Table). To determine juvenile and adult traits at the cellular level, we examined epidermal peels stained with Toluidine Blue O (TBO) [7]. Based on observations of bulliform cells, epidermal hairs, cell wall invagination, and TBO staining, all epidermal peels were juvenile at leaves 2-3 and began transitioning at leaf 4 (S7 Fig). NTG leaves were fully adult by leaf 7 and above, whereas UBI_pro_:*ZmCCT10* and BSV_pro_:*ZmCCT10* leaves were fully adult by leaf 8. There were no significant differences between NTG and transgenic plants for the juvenile-to-adult transition based on the cell morphology and TBO staining of epidermal peels. Therefore, we conclude that juvenile-to-adult phase change was not affected in these transgenic plants.

### Modification of reproductive traits in T1 transgenic plants

Transgenic plants exhibited a modification of reproductive traits. Relative to NTG plants, all transgenic plants displayed a delayed flowering phenotype (Table 2). NTG plants shed pollen on average 35-36 days after sowing. The UBI_pro_:*ZmCCT10* plants shed pollen ∼6 days later. Technical difficulties associated with maintaining the exceeding tall T1 BSV_pro_:*ZmCCT10* plants resulted in their final harvest and dissection at 130 days after sowing. On the day of harvest, only 51% (35 plants) of the BSV_pro_:*ZmCCT10* plants had shed pollen and they shed pollen 65 days later than NTG plants (Table 2), 43% (30 plants) produced the abnormal bushy tops, and 2 plants still had developing inflorescence meristem indicating an extreme delay in flowering.

**Table 2.** Reproductive traits collected for T1 families

| Line | Total No. Plants | Percent of plants that shed | Days to Shed | Percent of plants that silked | Days to Silk | Ear Length (cm) | Nodes with initiated primary ears | Shank Length (cm) | No. Primary Tassel Branches | No. Secondary Tassel Branches | No. plants with apically induced plantlets |
| --- | --- | --- | --- | --- | --- | --- | --- | --- | --- | --- | --- |
| NTG siblings | 60 | 100% | 36.2 ± 1.1 | 93% | 38.9 ± 2.7 | 9.2 ± 2.8 | 6 to 7 | 12.3 ± 6.0 | 9.0 ± 2.7 | 0 | 0 |
| UBI<sub>pro</sub>:*ZmCCT10* | 120 | 100% | 42.2 ± 4.4* | 100% | 48.8 ± 6.2* | 12.7 ± 1.8* | 6 to 8 | 31.2 ± 15.8** | 14.1 ± 3.2* | 0.7 ± 1.1 | 0 |
| NTG siblings | 45 | 100% | 35.6 ± 1.6 | 96% | 38.4 ± 2.5 | 8.6 ± 2.5 | 6 to 7 | 12.7 ± 5.4 | 7.2 ± 2.4 | 0 | 0 |
| BSV<sub>pro</sub>:*ZmCCT10* | 69 | 51% | 101.2 ± 12.9** | 16% | 94.3 ± 8.4** | 10.5 ± 2.9* | 18 to 33 | 13.3 ± 7.8 | 27.6 ± 7.2** | 10.9 ± 4.6** | 30 |

Measurements represent means ± SD. Only one BSV<sub>pro</sub>:*ZmCCT* event produced ears that silked (11 plants). \*Means are statistically different from NTG at p < 0.001. \*\*Means are statistically different from NTG at p < 0.00001.

Transgene expression had a significant effect on tassel branching. NTG plants produced tassels with an average of 7-9 primary branches and no secondary branches (Table 2 and S8 Fig). The UBI_pro_:*ZmCCT10* plants produced tassels with an average of 14 primary branches and, in a few examples, secondary branches were observed. The BSV_pro_:*ZmCCT10* plants developed highly branched tassels with an average of 28 primary and 11 secondary branches (Table 2 and S8 Fig).

For the NTG and UBI_pro_:*ZmCCT10* plants, 93-100% of the plants produced ears that silked 2 to 6 days later than they shed pollen. The unusually long shank subtending the ears observed in T0 UBI_pro_:*ZmCCT10* plants was reproduced in the T1 families. The average shank length of the UBI_pro_:*ZmCCT10* plants was 31 cm which was a 3-fold increase compared to the ∼12 cm long shanks produced by NTG, or BSV_pro_:*ZmCCT10* plants (Table 2). After careful dissection of the mature T1 plants we found that the BSV_pro_:*ZmCCT10* plants initiated ears but most them had aborted at early stages of development and never exerted silks. There was one event (11 plants) of the BSV_pro_:*ZmCCT10*_PI422162_ construct that produced silking ears. These plants were used to measure ear and shank length for the BSV_pro_:*ZmCCT10* plants (Table 2). The transgenic ears were ∼2 cm longer than NTG ears (Table 2).

### A high level of *ZmCCT10* expression caused formation of apically-induced plantlets

A striking feature of the BSV_pro_:*ZmCCT10* transgenic plants was the development of the “bushy top” phenotype upon plant maturity (Fig. 1D). More than 50% of the T0 plants developed this phenotype (S2E Fig), which was reproduced in 43% of the T1 BSV_pro_:*ZmCCT10* plants (Table 2). This feature was never observed in the NTG or UBI_pro_:*ZmCCT10* plants suggesting a strong relationship between the bushy top phenotype and the higher level of *ZmCCT10* transgene expression.

Dissection of the highly branched vegetative structure from the T0 plants revealed complex arrangements of vegetative structures resembling multiple plantlets, each with an individual shoot axis (Fig 3). The number of countable plantlets varied from 2 to 26 (S2C Fig). Plantlets were at different stages of development with larger, more mature plantlets positioned on the periphery of the tassel and younger, less developed plantlets closer to the center. Some apices still had a central growing point that could generate more plantlets (Fig 3B and 3C). Dissection of the individual plantlets revealed a complex mix of abnormal tassel-like and ear-like structures. In a few cases, ear-like structures were fertile and could be fertilized to produce kernels (Fig 3A, Row 4). Dissection of the T1 BSV_pro_:*ZmCCT10* bushy tops revealed the presence of apices consisting of only plantlets or a combination of plantlets and tassel branches (S9 Fig). The most mature looking plantlets (S 10A) when planted into soil developed large root systems (S10B, S10C and S10D Fig). However, the shoot growth of these transplanted plantlets was stunted with secondary plantlets developing in place of reproductive organs (S10C and S10D Figs).

**Fig 3.**
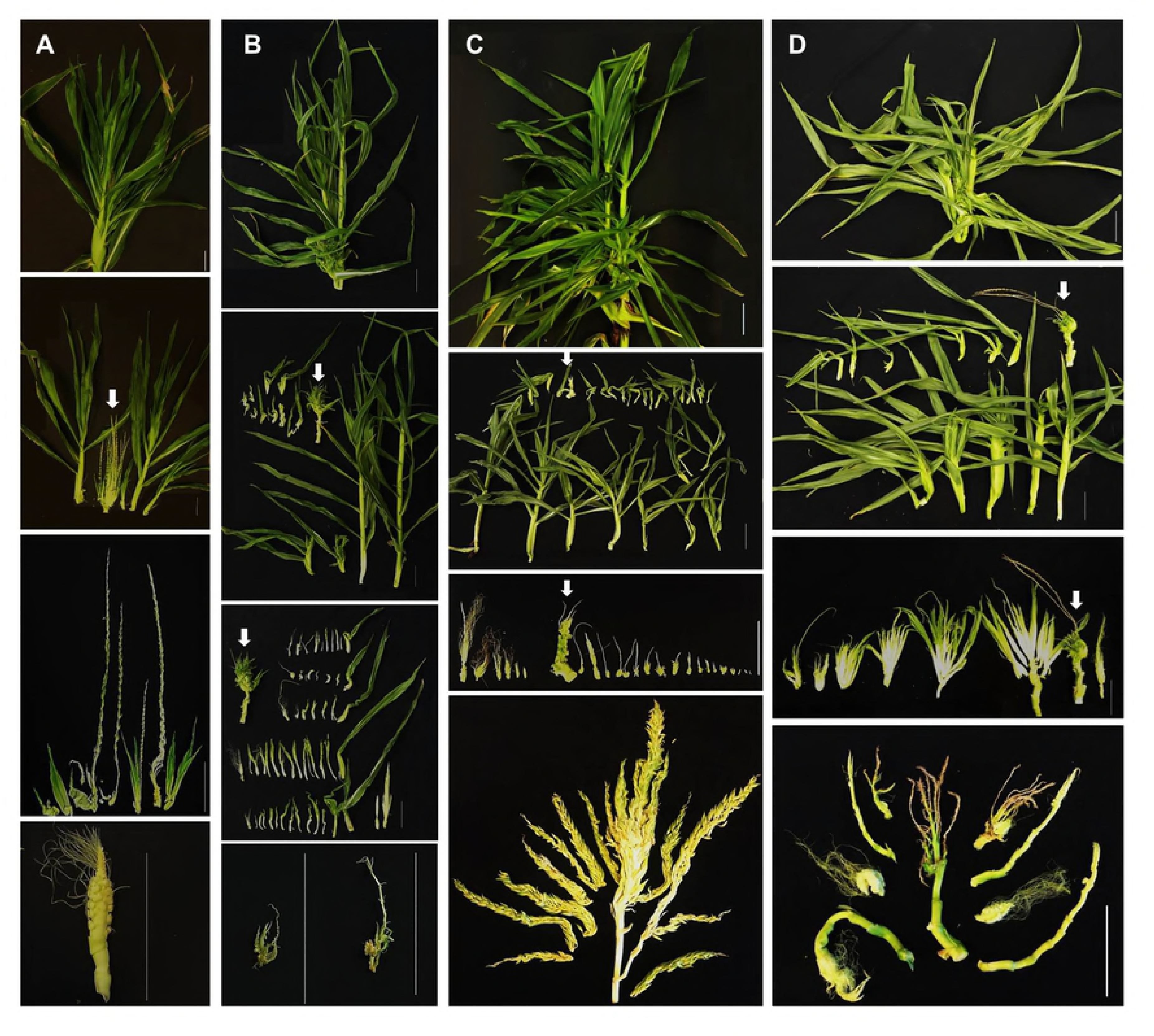
Representative images of apically-induced plantlets (bushy top) dissected from T0, BSVpro:*ZmCCT10* transgenic plants. Panel images are arranged in the following horizontal order - Row 1: Total view of apex of plant, Row 2: Individual apically-induced plantlets dissected from the apex above, Row 3: Internal structures of dissected apically-induced plantlets, Row 4: Internal structures of a single apically-induced plantlet showing propagation of phenotype (left) and additional dissection of the apex. Panel images are arranged in the following vertical order - A) BSVpro:*ZmCCT10*_*Gaspe*_ plants. B) BSVpro:*ZmCCT10*_B73_ plants. C) BSVpro:*ZmCCT10*_tropical_ plants. D) BSVpro:*ZmCCT10*_teosinte_ plants. White arrows indicate the central growing point. Scale bars = 10 cm.

To capture morphological changes in the meristem during and after floral transition, we dissected and imaged shoot apices from NTG and T1 transgenic plants. NTG plants developed very fast, as is typical for the Gaspé Flint line. Transitioning of the vegetative SAM to an inflorescence meristem (IM) occurred between V1-V2 stages (Fig 4A and 4B) followed by branch meristem (BM) initiation (Fig 4C) and developing tassel (DT) (Fig 4D) at V2-V3. By the V4 stage the immature tassel (IT) is fully formed and committed to maturation (Fig 4E). The apices of the UBI_pro_:*ZmCCT10* transgenic plants were sampled every V-stage until an immature tassel was formed (Fig 4F-J). The floral transition occurred at the V3-V5 stages, two stages later than NTG plants (Fig 4G). UBI_pro_:*ZmCCT10* apices showed initiation of multiple BMs (Fig 4H) that resulted in highly branched tassels (Fig 4I, 4J, and S8B). Due to the limited number of BSV_pro_:*ZmCCT10* T1 plants, we sampled apices at fewer V-stages and were only able to observe the vegetative SAM up to the V19 stage (Fig 4K). At this late stage, the SAM was swollen at the base (Fig 4K), that was not observed in NTG or UBI_pro_:*ZmCCT10* plants (Fig 4A and 4F). During removal and dissection of the very tall, latest flowering BSV_pro_:*ZmCCT10* plants at 130 days after sowing, we found two plants with an IM, and several plants with developing tassels (DT), which were characteristically highly branched (Fig 4L and 4M). Interestingly, two BSV_pro_:*ZmCCT10* plants were found that displayed a developing bushy top, with a developing tassel and plantlets initiated at lower positions (Fig 4N).

**Fig 4.**
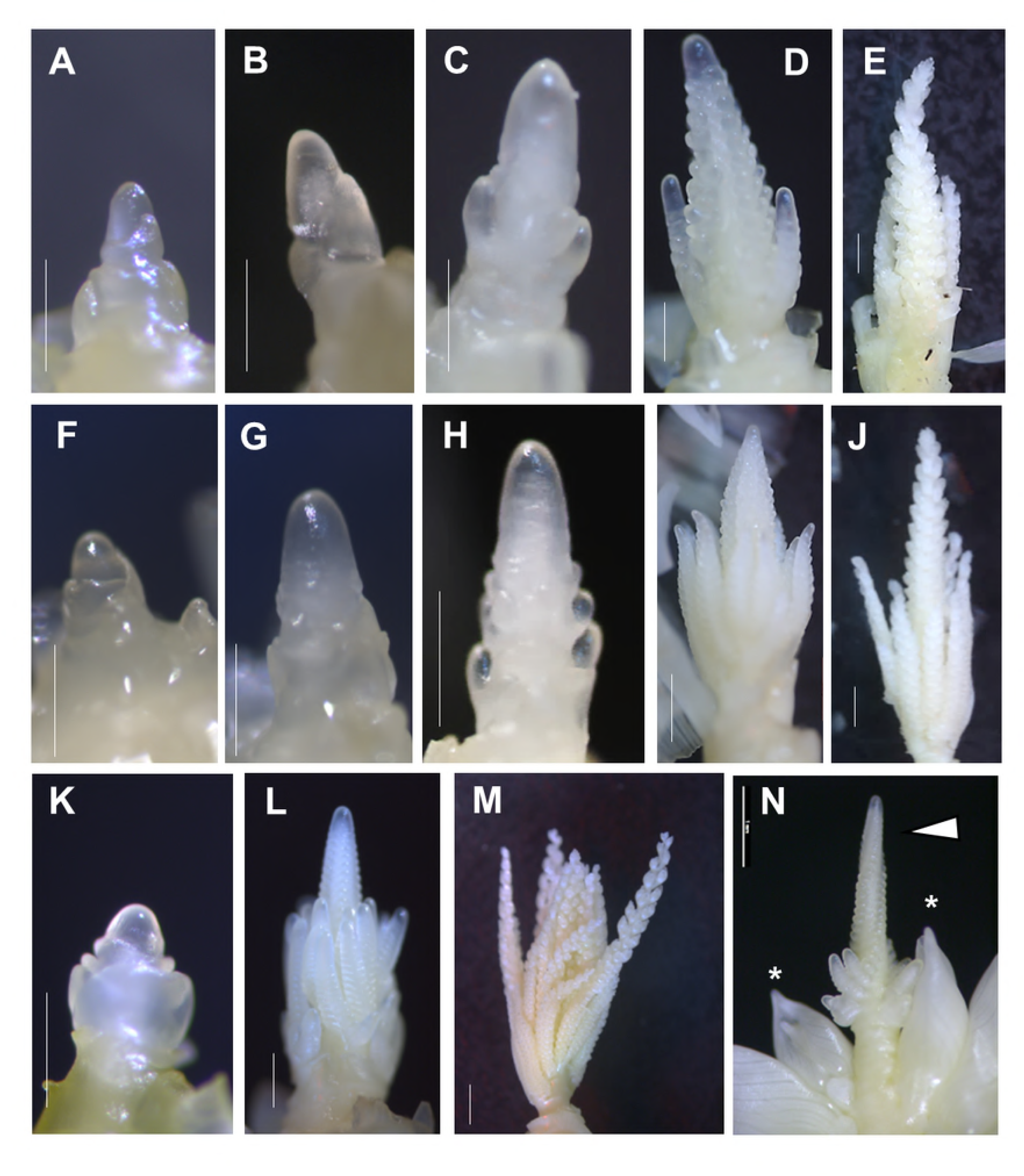
Representative images of apices dissected from non-transgenic and UBIpro:*ZmCCT10*, and BSVpro:*ZmCCT10* transgenic plants during vegetative and reproductive development. A to E) Apices of non-transgenic plants. F to J) Apices of UBIpro:*ZmCCT10* plants. K to N) Apices of BSVpro:*ZmCCT10* plants. A, F, K) Shoot apical meristem (SAM) at the vegetative stage. B, G) Inflorescence meristem (IM). C, H) Branch meristem (BM) initiation. D, I, L) Developing tassel (DT), the stage when all BM are initiated and branches continue to initiate spikelet pair (SPM) and spikelet meristems (SM). E, J, M) Immature tassel (IT), the stage of tassel growth when all spikelets are initiated and meristems are consumed. (N) The apex with combination of the tassel spike (arrowhead) and emerging plantlets (stars). Scale bars (A, B, C, F, G, H, K, N) = 500 mm, (D, E, I, J, L, M) = 1 mm.

To investigate the morphology of the apical meristem in the bushy top plantlets, we dissected the plantlets from several T1 BSV_pro_:*ZmCCT10* plants. The younger plantlets had a vegetative appearing SAM while others had developed a typical BSV_pro_:*ZmCCT10* highly branched immature tassel (S11A, B, E and F Figs). The shoot meristem of older plantlets had a mixed morphology, showing a combination of an immature main tassel rachis with numerous shoot-like structures basally positioned (S11C and D, S11M - O Figs), which we expected might form secondary plantlets. Other abnormalities were observed in the plantlet apical meristems, including ear-like structures surrounded by leaf-like primordia (S11J, S11K and S11L Figs) and a dissected spikelet containing both anthers and a cluster of pistillate florets (S11P, S11Q and S11R Figs).

### Expression of flowering regulators in leaves affected by *ZmCCT10* overexpression

To gain insight into the gene network regulated by *ZmCCT10* in leaves, we selected candidate genes based on the regulatory flowering network models proposed for maize [59] and rice [60, 61]. Even though *ZmCCT10* might regulate as many as 1117 genes in leaves [38], we selected 20 representative genes with proven or predicted functions in flowering pathways in leaves and assayed their expression by qRT-PCR (S5 Table).

The circadian clock plays a central role in regulation of photoperiodic flowering time in plants and so several clock genes were assayed. The maize genes *GIGZ1A/1B* and *CONZ1* are homologs of Arabidopsis flowering genes *GIGANTEA (GI)* and *CONSTANS (CO)* [62]. As previously shown, the duplicated genes *GIGZ1A and GIGZ1B* have identical expression patterns [62]. For this reason, *GIGZ1A/1B* were tested in the same assay (S5 Table). *GIGZ1A/1B* were not expressed in the coleoptiles but from leaf # 1 and onwards *GIGZ1A/1B* exhibited a typical diurnal expression pattern with an evening peak (Fig 5A). Hence *GIGZ1A/1B* genes appeared to be not regulated by *ZmCCT10* and likely functions upstream of *ZmCCT10* in the photoperiod pathway, as is the case in rice [60]. *CONZ1* gene expression was slightly reduced in the UBI_pro_:*ZmCCT10* transgenic plants and was significantly repressed in the BSV_pro_:*ZmCCT10* plants suggesting its regulation by *ZmCCT10* (Fig 5B).

**Fig 5.**
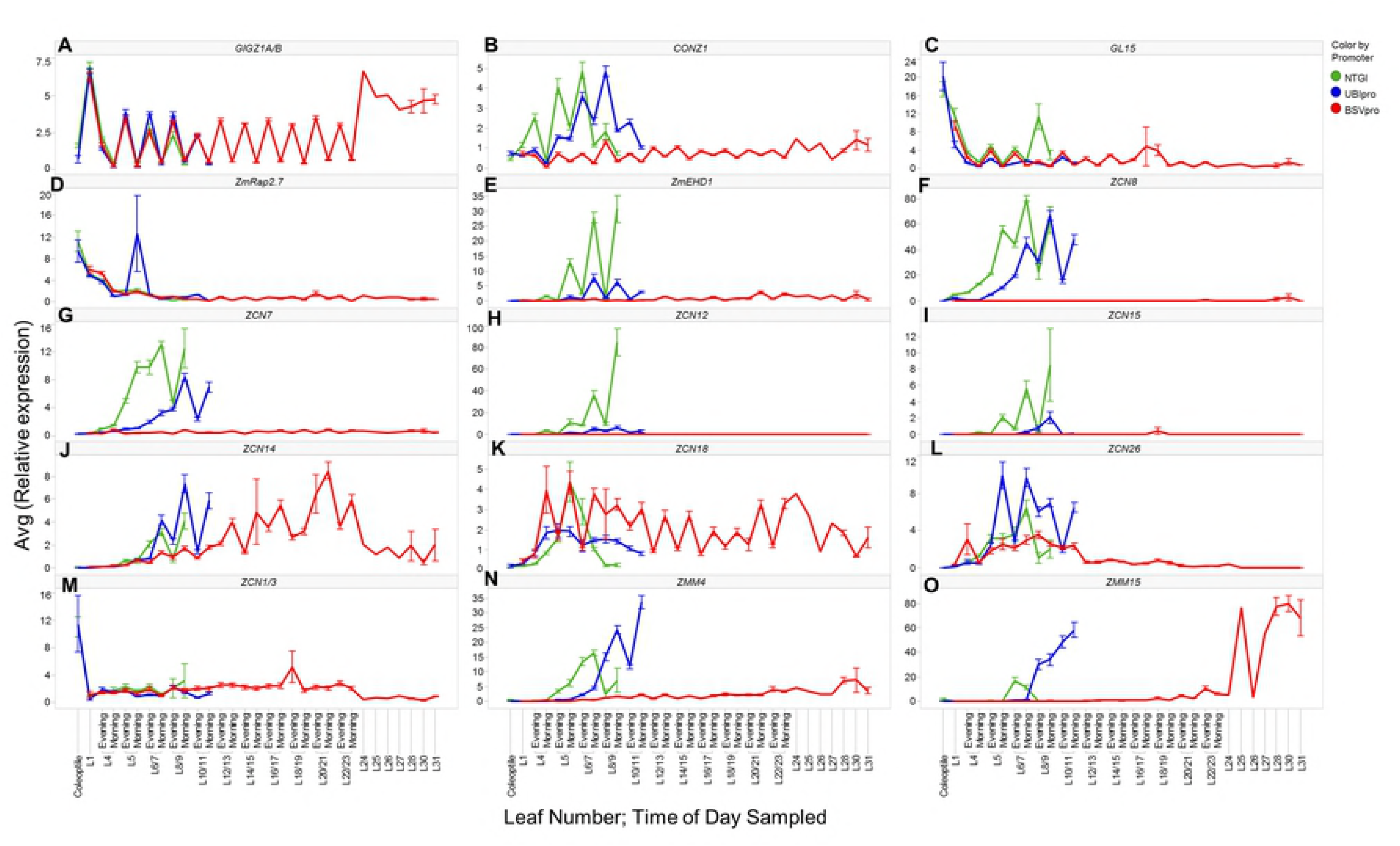
Expression patterns of flowering regulators in leaves of non-transgenic (NTG), UBIpro:Zm*CCT10*, and BSVpro:*ZmCCT10* transgenic plants. A) *GIGZ1A/1B, a* circadian clock gene. B) *CONZ1,* a circadian clock gene. C) *GL15*, the regulator of transition from a juvenile to adult leaf identity, a target of miR172. D) *ZmRap2.7,* a repressor of flowering, a target of miR172. E) *ZmEHD1,* an activator of flowering. F) *ZCN8*, the major florigen gene, an activator of flowering. G) *ZCN7*, the second florigen gene, an activator of flowering. H) *ZCN12,* I) *ZCN15,* J) *ZCN14,* K) *ZCN18*, FT-like genes with florigenic activity in Arabidopsis. L) *ZCN26, a* FT-like gene with no florigenic activity in Arabidopsis. M) *ZCN1/3*, TFL1-like genes, repressors of flowering. N) *ZMM4* and O) *ZMM15* MADS genes, activators of flowering, markers for floral transition. The X-axis represents the tissue sampled: Coleoptile, Leaves 1 to 22 (sampled either in the morning or evening), Leaves 24 to 31 (sampled in the evening). The Y-axis represents the average relative gene expression normalized against *eukaryotic initiation factor 4-gamma* (GenBank EU967723). Error bars represent ± the SE.

The *APETALA2*-*like* transcription factor *GL15 (GLOSSY15)*, that regulates the transition from juvenile to adult leaf identity in maize and is a target of miR172 [15], was highly expressed in coleoptiles and in juvenile leaf # 1. Thereafter, its expression decreased in leaf # 4 (Fig 5C), which is consistent with our observation that the onset of the transition from juvenile-to-adult phases occurred in leaf # 4 (S7 Fig). *GL15* was diurnally expressed with an evening peak and its expression pattern was nearly identical in NTG and transgenic plants suggesting no regulation by *ZmCCT10* (Fig 5C).

The *AP2-like ZmRap2.7* gene is a repressor of flowering and is a target of miR172 [63]. The *ZmRap2.7* expression pattern was similar to *GL15* which was higher in the coleoptiles and in juvenile leaf # 1 with a diurnal evening peak (Fig 5D). No significant differences were observed in NTG and transgenic plants placing *ZmRap2.7* outside of *ZmCCT10* regulation.

*ZmEhd1* is a homolog of the rice *Ehd1* activator of flowering (heading) under short days [42]. According to the rice photoperiod model, *Ehd1* is directly repressed by *Ghd7* [60, 61]. In agreement with the rice model, expression of *ZmEhd1* was reduced in the UBI_pro_:*ZmCCT10* transgenic plants and completely repressed in the BSVpro:*ZmCCT10* plants positioning *ZmEhd1* downstream of *ZmCCT10* (Fig 5E).

There are 15 *FT-like* genes in maize (the *ZCN* genes) that encode florigen-like proteins but only seven of them are expressed in leaves at some stages of development [28]. The *ZCN8* and *ZCN7* genes was shown to have florigen activity [29, 30, 64]. We examined the function of five leaf-expressed *ZCN* genes in Arabidopsis and found that four genes, *ZCN12/14/15/18,* displayed florigenic activity in Arabidopsis while *ZCN26* did not (S12 and S13 Fig). This finding suggested the potential for six ZCN proteins (including ZCN7/8) to function as florigen triggers in maize. Expression of all seven leaf-expressed *ZCN* genes was examined in the *ZmCCT10* transgenic plants. Expression levels of *ZCN8, ZCN7, ZCN12* and *ZCN15* were reduced in the UBI_pro_:*ZmCCT10* transgenic plants and completely repressed in the BSV_pro_:*ZmCCT10* transgenic plants (Fig 5F, 5G, 5H and 5I). Thus *ZCN8, ZCN7, ZCN12* and *ZCN15* appeared to be negatively regulated by *ZmCCT10*. Three of the *ZCN* genes, *ZCN14, ZCN18* and *ZCN26,* exhibited complex expression patterns (Fig 5J, 5K and 5L). *ZCN14* expression was slightly up-regulated in the UBI_pro_:*ZmCCT10* transgenic plants and its expression was continuously elevated in the BSV_pro_:*ZmCCT10* transgenic plants reaching its maximum in leaves 20-21 around the time when floral transition took place at least in some BSV_pro_:*ZmCCT10* plants (Fig 5J). Thus, *ZCN14* is not downstream of *ZmCCT10* and may potentially be influenced by long day-dependent mechanisms. *ZCN18* expression was slightly reduced in the UBI_pro_:*ZmCCT10* plants but was up-regulated in the BSV_pro_:*ZmCCT10* plants where it showed diurnal expression with a morning peak (Fig 5K). Thus, expression of *ZCN18* appears to be induced by high levels of *ZmCCT10* expression. *ZCN26* showed the opposite expression pattern of *ZCN18*. It was up-regulated in the UBI_pro_:*ZmCCT10* transgenic plants and was slightly repressed in the BSV_pro_:*ZmCCT10* plants in leaves #10-11 and completely repressed in older leaves (Fig 5L). Because *ZNC26* does not have florigenic activity and displays this unusual pattern, we speculate that its function may be unrelated to flowering time.

We also examined the *TFL1*-*like ZCN1/3* genes which are antagonists of *FT*-*like* genes and have been shown to delay flowering and modify plant architecture when over-expressed in maize plants [53]. Out of six *TFL1-like* genes (*ZCN1-6*), only *ZCN1* and *ZCN3,* are expressed in leaves at early growth stages [53]. The duplicated *ZCN1* and *ZCN3* genes showed higher expression in the coleoptiles and much lower in leaves at all stages of development in both NTG and transgenic plants (Fig 5M) and thus, are likely not regulated by *ZmCCT10*.

The *MADS* box genes, *ZMM4* and *ZMM15* are markers of the floral transition in maize. They are activated during this transition first in the reproductive inflorescences and then in the leaves [33]. *ZMM4* expression increased in leaves two stages later after the floral transition in NTG and UBI_pro_:*ZmCCT10* plants (Fig 5N). In the BSV_pro_:*ZmCCT10* plants, a slight expression increase was observed in leaves # 28-30 (Fig 5N). *ZMM15* expression increased in leaves # 6-7 of NTG plants and 2 leaf stages later in the UBI_pro_:*ZmCCT10* transgenic plants (Fig 5O). In the BSV_pro_:*ZmCCT10* transgenic plants, *ZMM15* expression was first detected in leaf #22 and increased in leaves # 24-31 reaching a higher level than in NTG or UBI_pro_:*ZmCCT10* plants (Fig 5O). This finding suggests that both *ZMM4* and *ZMM15* are regulated by *ZmCCT10* but *ZMM15* may be activated independently of the photoperiod pathway.

### Expression of meristem identity genes in the shoot apices affected by *ZmCCT10* overexpression

To gain insight into the gene network(s) regulated by *ZmCCT10* in the shoot apex, genes were selected with known and predicted functions in the floral transition and inflorescence development (S5 Table). For qRT-PCR analysis, the apices were grouped into five categories according to their developmental stage by morphology, including vegetative SAM, reproductive inflorescence meristem (IM), branch meristem (BM) initiation, developing tassel (DT), and immature tassel (IT) (Fig 4). Due to the limited number of BSV_pro_:*ZmCCT10* plants available for dissection, fewer apices from this genotype were sampled and those at the BM initiation stage were missed. Overall, we surveyed 30 genes with various meristem functions (S5 Table). Expressions of 20 genes were different in transgenic apices in more than one developmental stage (S6 Table). As a general trend, the magnitude of transcriptional changes was higher in the BSV_pro_:*ZmCCT10* apices compared to UBI_pro_:*ZmCCT10* apices (Fig 6 and S14).

**Fig 6.**
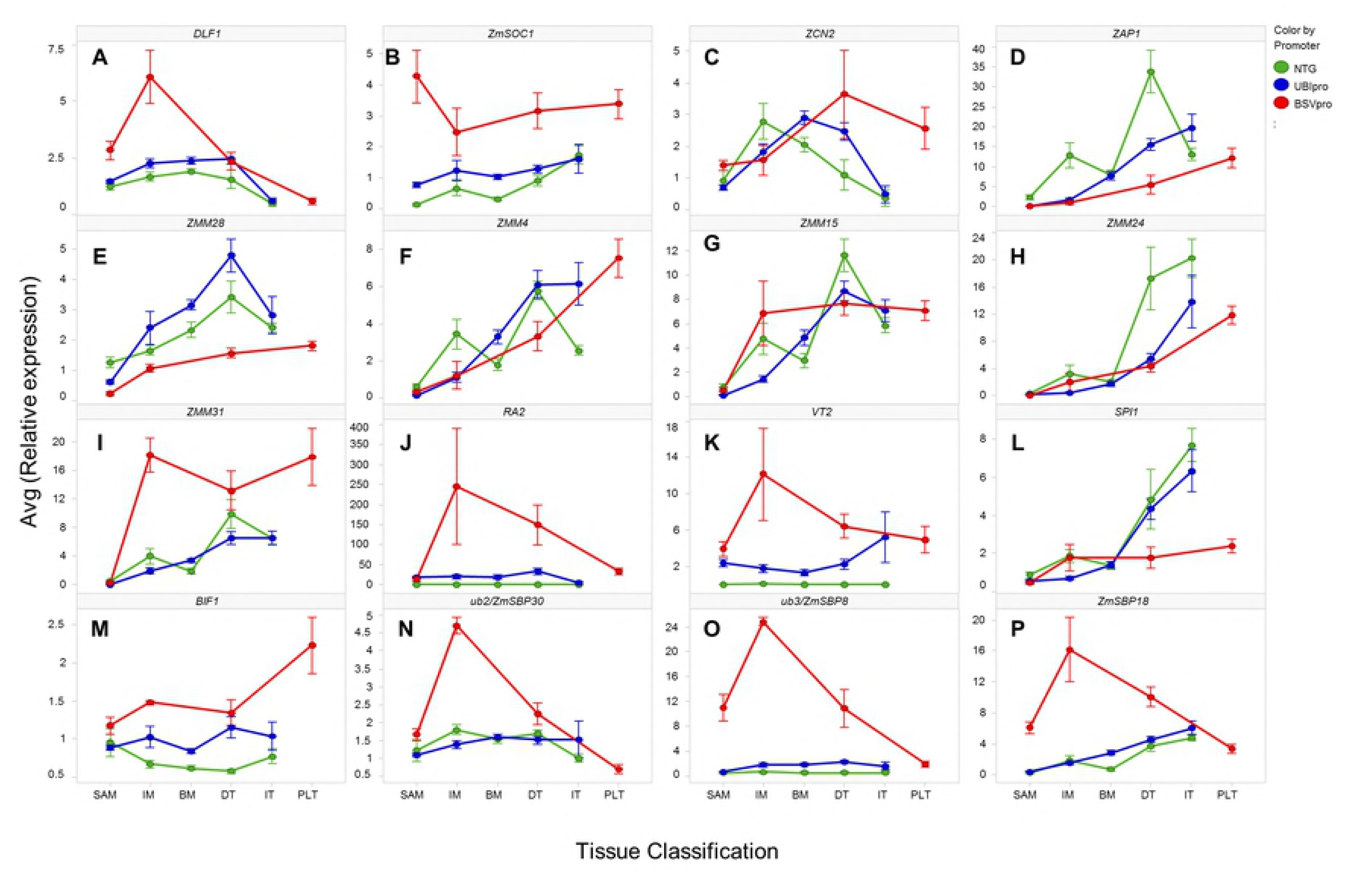
Expression patterns of inflorescence developmental genes in the shoot apices of non-transgenic (NTG), UBIpro:*ZmCCT10*, and BSVpro:*ZmCCT10* transgenic plants. A) *DLF1* and B) *ZmSOC1,* floral transition genes. C) *ZCN2,* maize homolog of Arabidopsis *TFL1,* a repressor of the floral transition. AP1-like MADS genes D) *ZAP1,* E) *ZMM28,* F) *ZMM4, and* G) *ZMM15*. SEP-like MADS genes H) *ZMM24* and I) *ZMM31.* J) RA2, SPM determinacy gene. K) *VT2* and L) *SPI1,* auxin biosynthesis genes. M) *BIF1,* auxin signaling gene. N) *UB2/SBP30. O*) *UB3/ZmSBP8, and P) ZmSBP18* genes, targets of miR156. The X-axis represents developmental stages of apices as defined in Figure 5. SAM: shoot apical meristem, IF: inflorescence meristem, BM: branch meristem, DT: developing tassel, IT: immature tassel, PLT: plantlets. The Y-axis represents the average relative gene expression normalized against *eukaryotic initiation factor 4-gamma* (GenBank EU967723). Error bars represent ± the SE.

The flowering activators *DLF1* [27] and *ZmSOC1* [65] known also as *ZmMADS1*[66] showed increased expression in transgenic apices (Fig 6A and 6B) suggesting their function independent of *ZmCCT10*. Expression of the flowering repressor *ZCN2*, a maize ortholog of Arabidopsis *TFL1*, showed similar expression trends in NTG and transgenic plants (Fig 6C). Expression of *ZFL1* that controls inflorescence phyllotaxy in maize [67, 68] was slightly lower in transgenic apices compared to NTG plants (S14A Fig, S6 Table) suggesting a partial regulation of *ZFL1* by the photoperiod pathway.

The inflorescence meristem identity *AP1*-*like* and *SEP*-*like MADS* genes were selected based on their key functions in inflorescence meristem specification in maize [33, 69] and rice [69-71]. As expected, the *AP1-like MADS* box genes, *ZAP1*, *ZMM28*, *ZMM4* and *ZMM15* were expressed at very low levels at vegetative stages in all genotypes. Their expression increased in the IM in NTG plants with *ZAP1* having the highest expression level (Fig 6D). In BSV_pro_:*ZmCCT10* apices, the expression levels of *ZAP1* and *ZMM28* were significantly reduced (Fig 6D and 6E) which indicates they function downstream of *ZmCCT10*. However, the expression of *ZMM4* and *ZMM15* was not significantly different in transgenic and NTG apices (Fig 6F and 6G).

The *SEP-like MADS* box genes *ZMM24* and *ZMM31* displayed contrasting expression patterns (Fig 6H and 6I). Expression of *ZMM24* was repressed in both UBI_pro_:*ZmCCT10* and BSV_pro_:*ZmCCT10*. But *ZMM31* expression was moderately increased in UBI_pro_:*ZmCCT10* apices at certain stages and was strongly increased in BSV_pro_:*ZmCCT10* apices. *ZMM31* expression was also high in plantlet apices (Fig 6I). This observation suggests that expression of the *AP1-like* and *SEP-like* genes is not tightly co-regulated by *ZmCCT10* even though AP1 and SEP MADS proteins seems to function together in specifying inflorescence meristem identity [71].

To address the morphological changes in the tassel, such as the increased number of branches, the *ramosa* (*RA*) genes that control branching were assayed [72, 73]. In NTG plants, *RA1, RA2, RA3* and *REL2* showed normal developmental expression patterns, consistent with original observations [73, 74]. Expression was not detected in the SAM, was detected in the IM, was increased in the BM and DT, and declined in the IT stages in NTG (Fig 6J, S14B, S14C and S14D). Although minor expression differences were observed for *RA1*, *RA3* and *REL2* in both UBI_pro_:*ZmCCT10* and BSV_pro_:*ZmCCT10* transgenic apices (S14B, S14C and S14D Fig), *RA2* expression was strongly increased in the transgenic plants, with BSV_pro_:*ZmCCT10* apices showing a 250-fold increase (Fig 6J). The *RA2* gene is required for transcriptional activation of *RA1* [75], thus one would expect *RA1* expression to also be up-regulated in transgenic plants, but this was not observed.

Given the critical function of auxin in plant development and shoot architecture [76, 77] we assayed auxin biosynthetic genes *VT2* [78] and *SPI1* [79]. Surprisingly expression of these genes showed opposing patterns (Fig 6K and 6L). Expression of *VT2* was significantly increased in UBI_pro_:*ZmCCT10* and BSV_pro_:*ZmCCT10* apices (Fig 6K) but expression of *SPI1* was either no different than NTG or was reduced in UBI_pro_:*ZmCCT10* and BSV_pro_:*ZmCCT10* apices (Fig 6L). Elevated expression of *VT2* was also observed in the roots of dissected apical plantlets (S10 Fig), but *SPI1* expression was not detected in roots. *VT2* encodes a tryptophan aminotransferase [78] and *SPI1* encodes a YUCCA-like flavin monooxygenase [79]. Both genes belong to different tryptophan dependent auxin biosynthesis pathways [79]. This finding suggests that those pathways may be regulated independently of each other and that regulation of the *SPI1* pathway might be influenced by the photoperiod. To further investigate effects on auxin pathway genes, expression of the auxin efflux transporter *ZmPIN1a* (*Zea mays pinformed1a*) [80] and the auxin signaling genes *BIF1, BIF4* and *BA1* [81] were assayed. Expression of *ZmPIN1a* was not affected in transgenic plants (S14L Fig). Expression of *BIF1* was increased for both transgenic constructs, whereas expression of *BIF4* and *BA1* were not statistically different (Fig 6M, S14M and S14N).

The novel phenotypic conversion of the tassel into a highly branched structure bearing plantlets in transgenic plants resembled the dominant *corngrass1* (*Cg1*) mutant which results from over-expression of miR156 and down-regulation of targeted *SBP* genes [12]. For this reason, we assayed six *SBP* genes regulated by miRNA156 cleavage including *TSH4/SBP6* (*tasselsheath4*) [82], *TGA1 (teosinte glume architecture1)* [12], *ub2 (unbranched2*) and *ub3 (unbranched3*) [83], and *ZmSBP14* and *ZmSBP18* [84]. None of six tested *SBP* genes showed decreased expression in transgenic plants compared to NTG plants (Fig 6N, 6O, 6P, and S14H, S14I and S14J Fig). In fact, increased transcript accumulation was observed for *ub2/ZmSBP30, ub3/ZmSBP8* and *ZmSBP18* (Fig 6N, 6O and 6P). Thus, it is likely that the apically induced plantlets are not caused by down-regulation of *SBP* gene expression.

To study the contribution of genes regulated by miR172, which has an antagonistic function to miR156, we selected the spikelet meristem identity genes *AP2*-*like* transcription factor *BD1 (branched silkless1)* [85], *TS6/IDS1 (tasselseed6/indeterminate spikelet1),* and *SID1 (sister of indeterminate spikelet1*) [86, 87]. The overall expression patterns of these genes were not statistically different between NTG and transgenic apices (S14E, S14F and S14G Fig), although *BD1* transcript accumulation was slightly decreased in UBI_pro_:*ZmCCT10* apices (S14F Fig).

We took advatage of the prolonged vegetative phase of transgenic plants to compare expression of the miR156 and miR172 targeted genes during aging of the SAM. The SAM of the UBI_pro_:*ZmCCT10* plants transtioned to an IM on average 20 days after sowing whereas the SAM of the BSV_pro_:*ZmCCT10* plants took up to 60 days. Transcript accumulation of the miR156-targeted genes, *ZmSBP18, TSH4/SBP6*, *TGA1, ub2/ZmSBP30, ub3/ZmSBP8* and *ZmSBP14,* showed an age dependent increase in expression (S15A, S15B, S15C, S15D, S15E and S15F Fig). This finding suggests that the expression of miR156 decreased over the extended vegetative period of growth resulting in accumulation of targeted *SBP* transcripts. Interestingly *TGA1* mRNA accumulation peaked around the floral transition in the apices of UBI_pro_:*ZmCCT10* and BSV_pro_:*ZmCCT10* transgenic plants (S15C Fig) suggesting a putative involvement in the transition from vegetative to reproductive development.

Transcript accumulation for the miR172-targeted *AP2-like* genes showed an inverse pattern to the *SBP* genes, with expression being higher in the younger stage SAM and declining in older SAMs (S15G, S15H, S15I Fig). This pattern suggests that expression of miR172 is low in young apices and increased during extended vegetative growth. This finding is consistent with observations that miR156-miR172 regulation of the juvenile-to-adult transition in leaves is not altered in transgenic plants and these miRNA species are not regulated by *ZmCCT10*.

## Discussion

### Duration of the vegetative phase shapes maize plant architecture

Genetic evidence suggests that *ZmCCT10* is a regulator of the photoperiod pathway in maize repressing flowering under long days [38, 48]. *ZmCCT10* is a nuclear localized protein with the potential to repress transcription, as was demonstrated for the homologous rice GHD7 protein [41]. For these reasons, over-expression of the ZmCCT10 protein in transgenic maize is expected to inactivate the photoperiod pathway and create plants mimicking growth under permanent long days. This transgenic model allowed us to study the impact of modulating the photoperiod pathway on plant growth and development as well as deciphering which genes connect to the photoperiod pathway in maize.

Over-expression of *ZmCCT10* with promoters of different strength, UBI_pro_:*ZmCCT10* and BSV_pro_:*ZmCCT10*, revealed that the phenotypic effects of *ZmCCT10* are quantitatively related to transcript abundance. Both UBI_pro_:*ZmCCT10* and BSV_pro_:*ZmCCT10* transgenic plants showed multiple changes to plant architecture, but the BSV_pro_:*ZmCCT10* transgenic plants developed the most extreme phenotypes. Quantitative transgene action may be explained by the diurnal turnover of the ZmCCT10 protein. This idea is supported by the observation that the homologous GHD7 protein in rice is degraded at night, but high protein levels accumulate each day correlated with high levels of *GHD7* transcription [41]. If this is the case for the ZmCCT10 protein, a higher transcriptional level may maintain a higher protein level.

Manipulation of expression of the *ZmCCT10* repressor with promoters of different strength revealed a strong relationship between duration of vegetative growth and plant architecture. As expected during the prolonged vegetative phase, plants produced more phytomers, recorded as the number of nodes with an attached leaf. This is clearly demonstrated by comparison of NTG Gaspé Flint plants with the shortest vegetative stage (1-2 days) producing a maximum of 10 leaves at maturity compared to the BSV_pro_:*ZmCCT10* transgenic plants with the longest vegetative stage (up to 60 days) producing up to 60 leaves. Since NTG Gaspé Flint plants transitioned to reproductive development at the V1 stage, when they are still in the juvenile phase, this suggests that phase change might not be required for initiation of reproductive development in maize.

Morphological parameters of the individual phytomer are also modified in transgenic plants. Internode length is shorter and stalk diameter is wider compared to NTG plants. Leaf shape along the shoot is also modified (Fig 2) with leaves above the ear continuing growth resulting in long, wide leaves on the upper shoot of transgenic plants (Fig 1). Transgenic BSV_pro_:*ZmCCT10* plants also developed adventitious brace roots at every internode up to the top of the plants (Fig 1). Another irregular phenotype produced only by UBI_pro_:*ZmCCT10* transgenic plants was an unusually long axillary branch (shank) subtending the ears (Fig 1D, Table 2). Extreme elongation of ear shanks was reported for the jasmonic acid-deficient double *opr7/opr8* mutants [88] hinting to a possible perturbation of jasmonic acid biosynthesis in UBI_pro_:*ZmCCT10* transgenic plants. All these modifications dramatically changed the overall architecture of the BSV_pro_:*ZmCCT10* transgenic plants indicating that prolonged vegetative growth could disturb multiple developmental processes.

Delaying the transition to reproductive development also induced changes in reproductive traits. We observed increased tassel branching in both UBI_pro_:*ZmCCT10* and BSV_pro_:*ZmCCT10* transgenic plants (S8 Fig) relative to the control. In BSV_pro_:*ZmCCT10* transgenic plants, all ears aborted, suggesting axillary inflorescence development was suppressed by increased apical dominance. Alternatively, constant strong *ZmCCT10* transcription may repress transition of the axillary branch meristem from vegetative to reproductive growth. It is important to emphasize that all maize plants with delayed floral transition display similar modifications to plant architecture. Production of more and bigger leaves, thicker stalks, brace roots at higher nodes, and highly branched tassels, was seen in the late flowering mutants *id1* [32] and *dlf1* [27]; in transgenic lines with less florigenic *ZCN8* gene activity [89], *TFL1-like* over-expression lines [53, 89], and tropically adapted maize lines grown under long days [36, 90]. Moreover, transformation of the tassel into a bushy top phenotype observed in the BSV_pro_:*ZmCCT10* transgenic lines (Fig 1), was similar to “a ball of shoots” phenotype observed in the null *id1-m1* mutant [32] and in some tropical lines grown under long day conditions [90]. Regardless of the genetic or environmental cues, prolonged vegetative growth leads to similar phenotypic modifications to plant architecture suggesting a common physiological mechanism. The nature of this mechanism remains unclear.

### Suppression of the photoperiod pathway does not affect transition from juvenile to adult phase but does delay the transition from vegetative to reproductive development

We found no effect of transgene expression on juvenile-to-adult phase change in transgenic events using such leaf morphological features as bulliform cells, epidermal hairs, cell wall invagination, and TBO staining (Fig S7). In agreement with phenotypic observations, expression of the *GL15* gene, a regulator of juvenile-to-adult phase leaf traits and a target of miR172 [15], is not affected in transgenic plants (Fig 5C). mRNA levels of miR172 targeted gene, *ZmRap2.7*, a repressor of flowering [63] also did not change in transgenic plants (Fig 5D). This finding suggests that the “adult” miR172-mediated aging pathway is not regulated by *ZmCCT10* and is not connected to the photoperiod pathway.

Higher *ZmCCT10* expression did result in a significant increase in leaf number and later time to flower which is evidence of repression of floral transition. We investigated the morphology of shoot apices and found that *ZmCCT10* expression delayed the transition of the vegetative SAM to an IM (Fig 4). The floral transition occurred two stages later in the UBI_pro_:*ZmCCT10* transgenic plants (V3-V4) compared to NTG controls (V1) and up to 20 stages later in the BSV_pro_:*ZmCCT10* transgenic plants. Suppression of the photoperiod pathway delayed transition from vegetative to reproductive development but had no effect on phase change.

### Disruption of meristem identity gene expression may cause the perturbed inflorescence morphology in *ZmCCT10* transgenic plants

We surveyed the expression of 30 meristem identity genes and found that for most of them, their expression was altered in BSV_pro_:*ZmCCT10* apices. The most common expression change was increased expression in the IM samples (Fig 6I, 6J, 6K,6N, and 6P). Among them, the two most striking were the 250-fold increase in *RA2* expression (Fig 6J) and the 12-fold increase in expression of the auxin biosynthesis gene *VT2* (Fig 6K), suggesting that auxin levels might be elevated in transgenic apices that contribute to alteration of meristem development. The mechanisms by which over accumulation of *RA2* or *VT2* contributes to the highly branched, vegetative bushy top phenotype is not clear. In fact, *RA2* normally functions to suppress the initiation of long branches on the tassel, since loss of function *ra2* mutants have highly branched tassels. Perhaps, increased branching in the *ZmCCT10* transgenics is mediated by a different branching pathway, like the *ub2/ub3* pathway, and *RA2* expression is activated in response to suppress the extra branching. Sorting out the cause and effect expression differences will require additional experimentation.

### *ZCN14* encodes a photoperiod independent long day florigen in maize

Tropical maize is a facultative long day plant [36] implicating the existence of an alternative pathway(s) to the photoperiod pathway that allows flowering under none-permissive long days. The photoperiod regulator *ZmCCT10*, when over-expressed in transgenic plants, represses expression of the two major florigenic genes *ZCN8* and *ZCN7* as well as the *FT-like* genes *ZCN12* and *ZCN15* (Fig 5F to 5I) placing them downstream of *ZmCCT10* in the photoperiod pathway. Expression of the *FT-like* genes *ZCN14* and *ZCN18* are not affected by *ZmCCT10* over-expression and thus, they function outside of the photoperiod pathway (Fig 5J and 5K). The expression of *ZCN14* in *ZmCCT10* transgenic plants is the same as its native expression in temperate and tropical lines under a long day photoperiod, supporting our original hypothesis that *ZCN14* promotes flowering under LDs [30]. The ZCN14 protein is phylogenetically related to the rice (Hd3a and RFT1), barley and wheat florigenic proteins [28] and interacts with DLF1 in a yeast two-hybrid system [30]. Therefore, we propose that the ZCN14-DLF1 complex forms a FAC under long day conditions (LD FAC) providing an alternative mechanism to flowering than the ZCN8-DLF1 FAC. It is important to emphasize that *DLF1* expression is not suppressed in BSV_pro_:*ZmCCT10* apices but is higher than in NTG and UBI_pro_:*ZmCCT10* plants (Fig 6A), indicating that the DLF1 protein is not rate limiting in forming the LD FAC.

The expression data from shoot apices suggest that the LD FAC may activate expression of the meristem identity genes *ZMM4 and ZMM15* because their expression is increased in BSV_pro_:*ZmCCT10* apices. Interestingly the *SEP-like MADS* box gene *ZMM31,* which is linked to *ZMM15* on the short arm of chromosome 5 [33], also showed an expression pattern similar to *ZMM15* in BSV_pro_:*ZmCCT10* apices. Thus, the alternative long day pathway seems to be composed of the genetic module *ZCN14* - *ZMM4* - *ZMM15* - *ZMM31*.

### A conceptual network model for regulation of flowering time in maize

A genetic network model for flowering time in maize [59] is less elaborated compared to rice [91] due to a limited number of flowering time mutants and a lack of flowering time QTLs with a large effect [92]. Transgenic manipulation of flowering time provides novel information with which to populate the maize network. Over-expression of *ZmCCT10* helped define the components of the photoperiod dependent and independent pathways and allowed us to further refine the flowering network (Fig 7).

**Fig 7.**
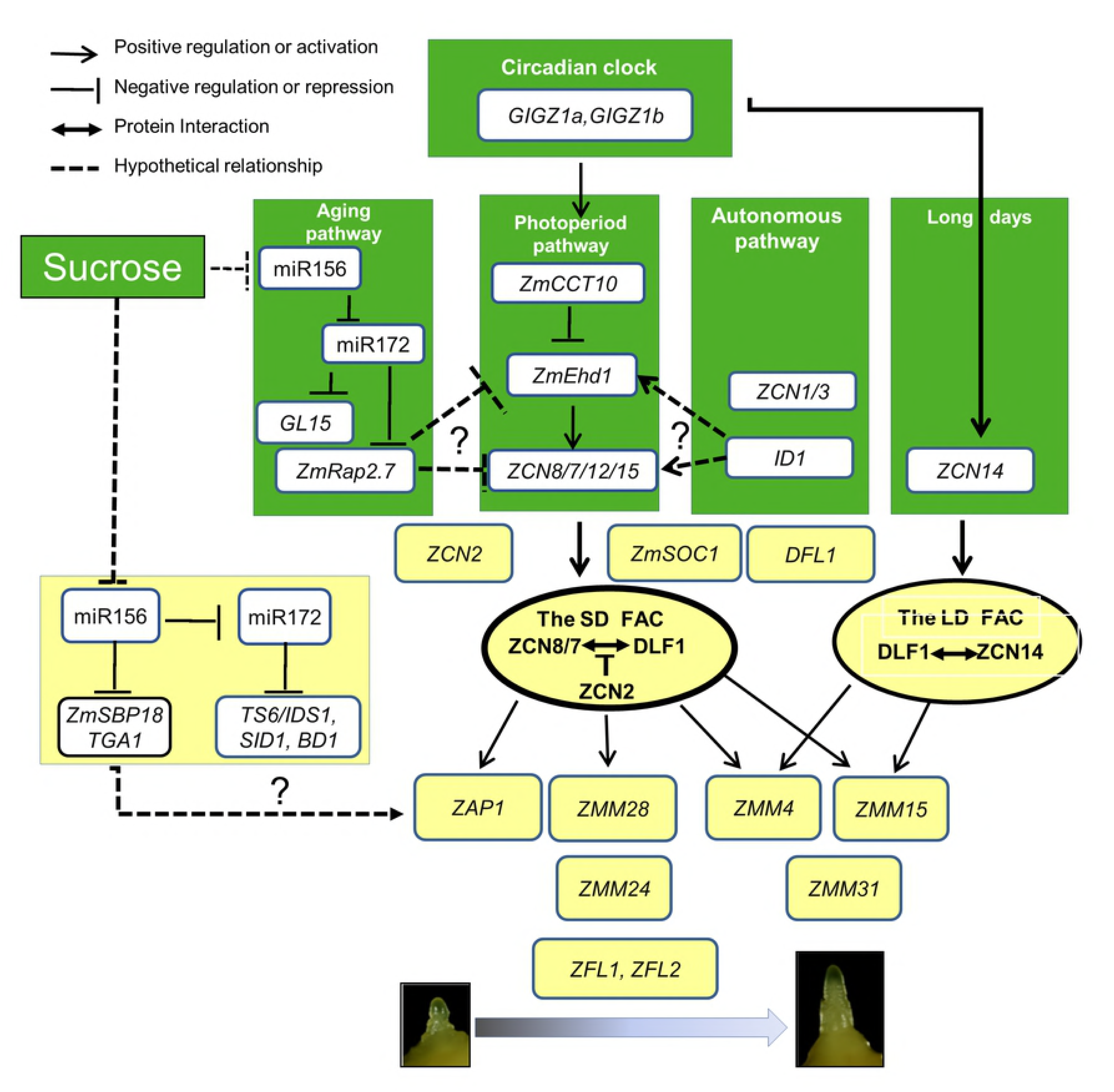
A conceptual gene network model for regulation of flowering time in maize. The model is separated by leaves (green background) and the shoot apical meristem (yellow background). In the leaves four pathways are depicted: aging, photoperiod, autonomous and long day. The circadian clock is the primary regulator of photoperiod and long day pathways. The photoperiod pathway is represented by the module *ZmCCT10* (rice *Ghd7*) - *ZmEhd1*– *ZCN8/7/12/15* (FT-like genes) which is conserved in other short-day monocots, rice and sorghum. *ZmCCT10* is a repressor of the flowering activator *ZmEhd1* gene. The second *ZmCCT9* might be positioned upstream of *ZCN8* but its precise position and relationship with *ZmCCT10* requires an additional study. The photoperiod independent long day pathway is represented by the florigenic gene *ZCN14* which expression is induced under the long days. The aging pathway *ZmRap2.7* gene acts as repressors of flowering that could act via repressing *ZmEhd1* or *ZCN8/7* genes. Expression of the autonomous pathway *ID1* gene is required for activation of *ZCN8/7* but directly or indirectly via activation of *ZmEhd1* remains to be determined. In the shoot apical meristem florigenic proteins ZCN8/7, transported from leaves, interact with the bZIP transcription factor DLF1 forming the florigen activation complex (the SD FAC). The antagonistic to florigen ZCN2 protein may compete with ZCN8/7 in the FAC leading to delayed flowering. Expression of *DFL1* and *ZCN2* is independent of the photoperiod pathway. Under the long day in tropical lines the LD FAC is formed by interaction of DLF1 and ZCN14 proteins. The SD FAC induced transcription of AP1-like MADS box meristem identity genes ZAP*1, ZMM4* and *ZMM15* marking the onset of inflorescence development. A target of the LD FAC is most likely *ZMM15*. Expression of the AP1-like genes demarcates the onset of the inflorescence development. The data does not provide enough resolution to position SEP-like MADS box genes relatively to the AP1-like genes. AP1-like and Sep-like MADS box transcription factors may form functional complexes. *ZFL1/2* genes are positioned the downstream of AP1-like genes. The aging pathways are regulated by sugar levels and some of *SBP* genes may provide the additional activation of the AP1-like genes which remains to be determined.

In the current network, the photoperiod pathway is represented by the conserved module of *ZmCCT10* (homolog of *Ghd7*) - *ZmEhd1* - *ZCN8/7* (FT-like florigen). Over expression of *ZmCCT10* does not affect expression of the circadian genes *GIGZ1A/B* (homolog of *GIGANTIA*), indicating the lack of feedback from *ZmCCT10* to the circadian clock. Expression of *ZmCCT10* might to be regulated by *GIGZ1A/B* similar to rice and sorghum. Maize *gigz1* mutants do not affect flowering under SDs but flowered slightly early under LDs in the day neutral background [93]. The *conz1* gene, the maize homolog of *CONSTANS,* is down regulated in *ZmCCT10* transgenic plants but its connection to other genes in this pathway is not clear.

In temperate day neutral lines, expression of *ZmCCT10* is disrupted by insertion of a transposon in the promoter region releasing the flowering activator *ZmEhd1* from its repressive action [38]. In rice, *Ehd1* is the upstream activator of the florigenic genes *Hd3a/RFT1* and whose expression is controlled by multiple upstream genes [91, 94]. In maize, *ZmEhd1* is likely to play a similar critical activation function of florigenic genes but this remains to be determined.

In *ZmCCT10* transgenic plants, expression of the florigenic genes *ZCN8/7/12/15* is repressed, positioning them downstream of *ZmCCT10*. In contrast, expression of *ZCN14* increases over time in *ZmCCT10* transgenic plants, suggesting it functions in the long day pathway. In day neutral plants, *ZCN14* expression is observed after the floral transition in leaves, ovules and kernels [28] suggesting it may have a novel function in the absence of selection under long days.

In the shoot apical meristem, *DLF1* is expressed independent of the photoperiod pathway, allowing for a constant supply of this bZIP transcription factor to the FAC that was first described in rice [24]. In day neutral and tropical lines under SDs, the primary florigen is supplied by *ZCN8* gene which is transported to the SAM [29, 30] where it interacts with DLF1 to form the FAC. This complex activates expression of the meristem identity AP1-like MADS box genes *ZAP1, ZMM28, ZMM4* and *ZMM15*. An alternative flowering pathway under LDs may function through a long day FAC (LD FAC) that is formed by DFL1 and ZCN14 which seems to activate only *ZMM4* and *ZMM15* meristem identity genes.

The aging pathways in leaves and apical meristems are not repressed in *ZmCCT10* transgenic plants which places them outside of the photoperiod pathway. However, during prolonged vegetative growth, we observed age-dependent activation of *SBP* genes in the SAM. This finding suggests that some SBP proteins might have the ability to bypass the need for FAC function and activate *AP1-like MADS* box genes independent of the florigen-dependent pathway for flowering. A similar situation was shown to function in Arabidopsis where expression of the *spl3* and *spl9* (*squamosa promoter binding protein-like*) genes could trigger expression of *AP1, FUL1 and LEAFY* and, thus, bypass the need for FT-FD function to induce the floral transition [95, 96]. If a similar bypass mechanism functions in maize it has yet to be determined.

## Conclusion

Maize was domesticated from the tropical progenitor-grass teosinte, which requires short-day photoperiods to flower. Over time, as cultivation moved to higher latitudes, maize’s requirement for short-day photoperiod induction was reduced. As a result, maize has been adapted to grow in a wide range of photoperiods, with tropical maize requiring short days to initiate reproductive development, and temperate maize being relatively day neutral. Genetic studies revealed that *ZmCCT10*, which functions as a repressor of flowering, controls this short-day requirement. We investigated the role this gene plays in flowering and other developmental processes by generating transgenic plants overexpressing *ZmCCT10*. Our phenotypic analysis of transgenic events containing either a constitutive promoter with high strength, or a constitutive promoter with very-high strength, showed that *ZmCCT10* over-expression produced the expected late flowering phenotype, with the very-high level expressing events showing a dramatically prolonged vegetative period of growth and severe morphological developmental defects. Transcript expression analyses indicated that many genes that promote flowering are repressed and thus are downstream of *ZmCCT10*. We also showed that specific genes in modules affecting other developmental transitions are linked to photoperiod dependent or independent regulation. Our analysis allowed to propose an updated conceptual flowering time model for maize.

## ACKNOWLEDGMENTS

We are thankful to Ron Christensen, Jacque Hockenson and Tim Moriarty for assistance with growing plants in the green house; Kevin Simcox for supplying the seeds of teosinte, Shawn Thatcher for help with the statistical analysis and Gene Ananiev for figure preparation.

## Supporting information

**S1 Fig. Multiple alignments of ZmCCT10 amino acid sequences deduced from temperate Gaspé Flint and B73 lines, tropical CML311 and CML436 lines, wild progenitor teosinte (accessions PI 441934 and PI422162) and sorghum Sb-GHD7.** Amino acids conserved in all genotypes are shown over yellow background. Amino acids found in only one genotype are shown over green background. Amino acids conserved in five or six genotypes out of seven are shown over the blue background. The conserved CCT domain is framed. Asterisks marked putative DNA/RNA binding motif C-X2-C-X4-CC-X-H-X2-H. Putative nuclear localization signals (NLS) are underlined.

**S2 Fig. Quantification of modified traits in T0, BSVpro:*ZmCCT10* transgenic plants and non-transgenic plants (NTG) by allele.** NTG parent: no overexpression of *ZmCCT10* allele, Gaspe: overexpression of Gaspé Flint *ZmCCT10* allele, B73: overexpression of B73 *ZmCCT10* allele, CML436: overexpression of the CML436 *ZmCCT10* allele, CML311: overexpression of the CML311 *ZmCCT10* allele, PI422162: overexpression of the teosinte PI422162 *ZmCCT10* allele, PI441934: overexpression of the teosinte PI441934 *ZmCCT10* allele. A) Internode length by leaf position. Internode distance between nodes 4 and 5 is referred to as node 5. Measurements represent means ± SD. B) The range of the highest nodes with the brace roots depicted by the box-plot. C) The number of apically–induced plantlets in T0 plants depicted by the box-plot. D) Example of secondary aerial brace roots formed at 153 days after planting. E) The percentage of T0 plants with normal and modified apex morphology “bushy top” by *ZmCCT10* alleles. Apex morphology is classified as shoot apical meristem (SAM) in the vegetative state, the tassel, both tassel and plantlets, as well as plantlets only.

**S3 Fig. A BSVpro:*ZmCCT10* transgenic stalk withstands 80°bending.**

**S4 Fig. Relationship between specific traits and the level of *ZmCCT10* transgene expression in T1, non-transgenic (NTG), UBIpro:*ZmCCT10*, and BSVpro:*ZmCCT10* transgenic plants.** A) Plant height at harvest. B) Final leaf number. C) The highest nodes with brace root initiation. D) The highest nodes with primary ears (aborted ears in the BSVpro:*ZmCCT10* transgenic plants).

**S5 Fig**. **Representative images of T1, non-transgenic plants and transgenic BSVpro:*ZmCCT10* plants focusing on the base of the plants.** Scale Bar = 1 m.

**S6 Fig. Linear regression analysis of leaf appearance rate in T1 non-transgenic (NTG), UBIpro:*ZmCCT10,* and BSVpro:*ZmCCT10* plants.** The plant leaf number was recorded twice a week. Linear regression lines show leaf appearance rate. b-value indicates average number of leaves appearing in one day. r2 indicates how well the data fit the trend line.

**S7 Fig**. **Adaxial epidermal peels of non-transgenic (NTG) and transgenic (UBIpro:*ZmCCT10* and BSVpro:*ZmCCT10*) leaves stained with toluidine blue O.** A) Leaf 2 from NTG, B) UBIpro:*ZmCCT10* and C) BSVpro:*ZmCCT10* plants representing the juvenile phase. Juvenile epidermal cells are elongated, stain violet, and possess wavy cell walls. D) Leaf 4 from NTG E) UBIpro:*ZmCCT10* and F) BSVpro:*ZmCCT10* plants in the transitioning stage showing a mixture of juvenile and adult traits. Macrohairs are visible, but files of bulliform cells are not formed yet. G) Leaf 7 from NTG, and leaf 8 H) UBIpro:*ZmCCT10* and I) BSVpro:*ZmCCT10* plants representing the adult phase. J) Graphic representation of leaf identity vs. leaf number. The epidermis is highly differentiated into aqua-staining cells with invaginated cell walls, files of purple bulliform cells with macrohairs. b, files of bulliform cells; m, macrohair. Scale bar = 500 mm

**S8 Fig. Representative images of non-transgenic and transgenic tassels.** A) Non-transgenic tassels, B) UBIpro:*ZmCCT10* tassels, and C) BSVpro:*ZmCCT10* tassels. Scale bar = 5 cm

**S9 Fig. Images of apically-induced plantlets dissected from T1, BSVpro:*ZmCCT10* transgenic plants.** A,B,C) Examples of the apices that produced only plantlets. D,E,F,G) Examples of the apices that produced plantlets and tassels. Scale bar = 5 cm.

**S10 Fig. Detached apically-induced plantlets replanted in soil.** A) Images of 9 plantlets and the main growing stalk dissected from one T1 BSVpro:*ZmCCT10* plant. The more developed plantlets #1, #2 and #3 were planted into pots on June 30, 2015 and grown in a greenhouse until August 19, 2015. Plantlet #2 died. B) View of plantlet #1 showing well-developed roots (close-up in insert). C) Dissection of plantlet #1 showing continuous production of secondary plantlets. D) View of plantlet #3 showing well-developed roots, developed ear (close-up in insert on the right side) and the main growing stalk producing secondary plantlets (close-up in insert on the left side). Scale Bars = 30 cm.

**S11 Fig. Variations of impaired inflorescence development in apically induced plantlets in T1 BSVpro:*ZmCCT10* transgenic plants.** Plantlets dissected from the same plant are grouped and numbered starting from the most mature plantlets (#1). A), B), H) Visibly normal immature tassels. C), D) M), N), O) Apices with the developed main tassel spike (arrowheads) and emerging secondary plantlets (stars) at the base of the tassel. E), F) Apices with the vegetative SAM. G), I) Severely impaired tassels with leaf-like structures. J) Apices with massive outgrowth of ear-like structures at the base of the main tassel spike and emerging secondary plantlets (stars). K) A close-up view of ear-like structures. L) Dissection of the secondary plantlet from the apex [(J) marked by arrow] show the ear-like structure with tertiary emerging plantlets (stars). P) The staminate spikelet dissected from the tassel-like structure with 2 normal (instead of 3) stamens at the right-side floret, close-up Q) and the ear-like structure at the left side floret, closeup. R) The main tassel spikes are marked by arrowheads and emerging plantlets are marked by stars. Scale bars (A, B, K, L) = 1 mm, (G, H, I, O) = 2 mm, (J, M, N) = 5 mm, (C, D, E, F, P, Q, R) = 500 mm.

**S12 Fig. Phenotypes of T2 Arabidopsis *ft* mutants complemented with maize FT-like *ZCN* genes**. Scale bar = 2 cm.

**S13 Fig. Complementation of Arabidopsis *ft* mutants with maize FT-like *ZCN* genes.** Over-expression of *ZCN8* and *ZCN12* not only complements *ft* mutants but also causes extremely early flowering and determinate inflorescences. *ZCN12* T1 over-expression lines are sterile. Over-expression of *ZCN14*, *ZCN15*, and *ZCN18* complements *ft* mutants and causes early flowering and indeterminate inflorescences. Over-expression of *ZCN26* does not complement *ft* mutants and leads to a slightly late flowering. Number of rosette leaves are means ± SD.

**S14 Fig. Expression patterns of inflorescence developmental genes with no statistically significant differences in more than one stage in apices of non-transgenic (NTG), UBIpro:*ZmCCT10*, and BSVpro:*ZmCCT10* transgenic plants.** X-axis represents developmental stages of apices as defined in Figure 5. SAM: shoot apical meristem, IF: inflorescence meristem, BM: branch meristem, DT: developing tassel, IT: immature tassel, PLT: plantlets. Y-axis represents average relative gene expression normalized against *eukaryotic initiation factor 4-gamma* (GenBank EU967723). Numeric t-test, p>0.05 are shown in S4 Table where expression in the transgenic shoot apices was compared to NTG apices. Error bars = ± SE.

**S15 Fig. Age-dependent mRNA accumulation of *SBP* genes and *AP2*-like genes in the shoot apical meristems of UBIpro:*ZmCCT10* and BSVpro:*ZmCCT10* transgenic plants.** A) to F) Quantification of *SBP* mRNA, regulated by miR156 cleavage. G) to I) Quantification of *AP2*-like mRNA, regulated by miR172 cleavage. X-axis represents days after sowing (DAS). Y-axis represents relative gene expression normalized against *eukaryotic initiation factor 4-gamma* (GenBank EU967723). Each data point represents a qRT-PCR value for an individual apex.

**S1 Table. Phenotypic traits of T0 transgenic plants (UBIpro:*ZmCCT10* and BSVpro:*ZmCCT10*) separated by constructs.**

**S2 Table. Phenotypic traits of T1 transgenic plants (UBIpro:*ZmCCT10* and BSVpro:*ZmCCT10*) separated by constructs.**

**S3 Table. Stalk diameter at various nodes in non-transgenic (NTG) and transgenic (UBIpro:*ZmCCT10* and BSVpro:*ZmCCT10*) families.**

**S4 Table. Percent of plants with macro-hairs by leaf number in non-transgenic (NTG), UBIpro:*ZmCCT10*, and BSVpro:*ZmCCT10* transgenic plants.**

**S5 Table. Summary of genes analyzed from the flowering and meristem identity pathways.**

**S6 Table. T-test results of gene expression (qRT-PCR) in non-transgenic vs. transgenic plants (UBIpro:*ZmCCT10* and BSVpro:*ZmCCT10*) apices.**

